# The Case for Kinases: A Phosphorylation Driven Model for Circadian Temperature Compensation

**DOI:** 10.64898/2026.05.07.723636

**Authors:** Elizabeth-Lauren Stevenson, Christina M. Kelliher, Arminja N. Kettenbach, Jennifer J. Loros, Jay C. Dunlap

**Affiliations:** Dept. of Molecular and Systems Biology, Geisel School of Medicine at Dartmouth, Hanover, NH; Dept. of Biology, University of Massachusetts Boston, Boston, MA; Dept. of Biochemistry and Cell Biology, Geisel School of Medicine at Dartmouth, Hanover, NH

## Abstract

Circadian rhythms, ∼24-hour biological cycles, enable organisms to anticipate rhythmic environmental cycles so they can assign proper day and night functions that align with those cycles. Circadian rhythms are defined by their ability to be reset by external cues, their capacity to continue to oscillate in the absence of those cues, and their capacity to maintain the rate of the clock across a range of ambient temperatures, a property known as temperature compensation. In the *Neurospora* clock, the White Collar Complex (WCC) drives expression of FRQ which nucleates a complex including FRH and CK1a that phosphorylates and thereby represses WCC activity. Work to date has suggested that kinases may be involved in temperature compensation and that in *Neurospora* the primary target of these is FRQ. Here we investigate the genetic relationship between two clock kinases, Casein Kinase I (*ck-1a*) and Casein Kinase II (*cka*), in their regulation of temperature compensation using novel alleles, *ck-1a^D135G^* and Δ*cka*. We find that that the clock relies on Casein Kinase I more at cold temperature, but this changes as temperature increases, and the clock relies more on Casein Kinase II at warm temperatures. Using quantitative proteomics on FRQ across temperatures, we find that the FRQ phosphorylation landscape is dependent on temperature and is altered in temperature compensation mutants. This leads to the development of a phosphorylation driven model for temperature compensation, where key temperature compensation specific domains on FRQ are phosphorylated to regulate period length in response to temperature, including by Casein Kinase I and Casein Kinase II.

## Introduction

Circadian rhythms have evolved (at least three times^1–4)^ to enable organisms to anticipate daily environmental cycles. To accurately perform this function, the circadian oscillator must be able to maintain its rate (period length) across a range of environmental conditions^5^ that may otherwise impact the molecular reactions that underlie it, such as differences in temperature^6^. This capability, “compensation”, is a defining circadian principle^7,8^ and is therefore present in all circadian systems: the clock does not exist without it. The molecular mechanism underlying this essential property, however, has remained elusive. Several general models have been proposed to explain temperature compensation: 1) multiple temperature dependent processes that have opposing effects on the clock may balance one another out to maintain consistent period length across temperature^7,9^; 2) viewing the clock as a limit cycle, as temperature and therefore rate increases, if the size of the limit cycle (circadian amplitude) increases, the oscillation would take the same amount of time^10–12;^ and 3) the salient period determining step of the clock may be temperature compensated within itself^13–16^.

In animals and fungi, the core clock is comprised of a functionally conserved negative transcription translation feedback loop consisting of a heterodimer of transcription factors (the “positive arm”) that activate expression of core clock genes which form a complex and, in turn, feedback to inhibit the action of the positive arm and thereby their own transcription (the “negative arm”). This inhibition is accomplished in part through phosphorylation of the positive arm heterodimer by Casein Kinase I that is part of the negative arm complex, in addition to auxiliary kinases. Positive arm repression is reprieved to enable the start of a new cycle when the negative arm is hyperphosphorylated^17,18^, by action of Casein Kinase I and additional kinases^19^, or degraded^20^. In the filamentous fungus *Neurospora*, White-Collar 1 (WC-1) and White-Collar 2 (WC-2) make up the positive arm^21^, and Frequency (FRQ)^22^, FRQ-Interacting RNA Helicase (FRH)^23^, and Casein Kinase I (CKI)^24^ make up the negative arm. WC-1 and WC-2 perform the same function as the transcription factors BMAL1 and CLOCK in mammals, FRQ is functionally homologous to PER in mammals, and CKI is orthologous to CKIε/δ in mammals^25^. Because this regulatory architecture is conserved between *Neurospora* and mammals, insights gained into the molecular mechanism of the clock using the simple and genetically tractable fungus *Neurospora* have in nearly all cases translated into insights to how the clock functions in mammals^26^ as well.

In *Neurospora*, investigations into how temperature compensation may be maintained suggest that kinases lie at the heart of the mechanism. Casein Kinase II (CKII), a kinase known to phosphorylate FRQ^19^, was found to regulate temperature compensation with the identification of two known temperature compensation mutants, *period-3* (*prd-3*) and *chrono,* as hypomorphic SNPs in *cka* and *ckb-1,* respectively^27^, the genes the encode the catalytic and regulatory subunits of CKII. PRD-4, the ortholog of Checkpoint Kinase 2, causes under-compensation in *Neurospora* when hyperactive^28^. Several new lines of evidence suggest that CKI also contributes to temperature compensation regulation. CKI^H123Y^, which reduces CKI protein levels, was found to cause the clock to become under-compensated^16^. These and other data have given rise to a model which posits that compensation arises from maintenance of the FRQ-CK-1a interaction at a steady level across temperatures^16,29^. Indeed, CKIε/δ^30–32^ is also thought to be a primary regulator of temperature compensation in mammals^33^, as the CKIε^Tau^ mutation causes under-compensation in the mammalian clock^34,35^, and the compensation model in mammals relies on phosphorylation of PER by CKIε/δ. The current observations into the temperature compensation mechanism of other model organisms are reviewed elsewhere^36^.

Here we have used a variety of genetic tools to dig more deeply into the roles of kinases in temperature compensation distinct from period length determination. We generated a complete knockout of *cka*, the catalytic subunit of CKII, to show that the clock can oscillate (at a long period) without CKII, and that this knock-out is more over-compensated than the *cka* hypomorph, *prd-3*. We report a previously undescribed mutation, CKI^D135G^, that shows extreme under-compensation and find that reducing either CKI levels or activity also leads to under-compensation. We used quantitative phosphoproteomics to assess FRQ phosphorylation across temperature in WT and mutant backgrounds and find that FRQ phosphorylation *is* temperature dependent and is altered in strains with disrupted temperature compensation. We investigated the relationship between CKI and CKII in temperature compensation regulation through epistasis experiments using both genetic manipulation of CKI and the CKI-specific inhibitor PF-670462. We find that the genetic relationship between these kinases is nuanced, with CKI being the primary regulator of period length at cold temperatures, while CKII is the primary regulator of period length at warm temperatures.

### A complete knock-out of Casein Kinase II is overcompensated and its suppressor mutation, CKI^D135G^, is undercompensated

A previous report^27^ identified the over-compensated mutant, *prd-3*, as a hypomorph of CKII due to a SNP in *cka*, so we generated a homokaryon knock-out of *cka*, the catalytic subunit of CKII (**Fig. 1A**). We measured the period length of WT, *ras-1^bd^*, *ras-1^bd^ prd-3*, and Δ*cka::hph* at 20°, 25°, and 30°C using a luciferase clock reporter^17,37^ to determine the impact of these mutations on the clock’s ability to buffer against changes in ambient temperature. The *Δcka* mutant is viable and fertile as a male parent but displays very poor growth. We found that the complete knock-out of CKII has an even greater over-compensation defect than its corresponding hypomorph, *prd-3*, with an even more positive slope in its temperature compensation profile (**Fig.1A, S1A**). See Supplemental Information for a table of the Q10 coefficient calculations for all strains reported in this study.

**Figure 1.**
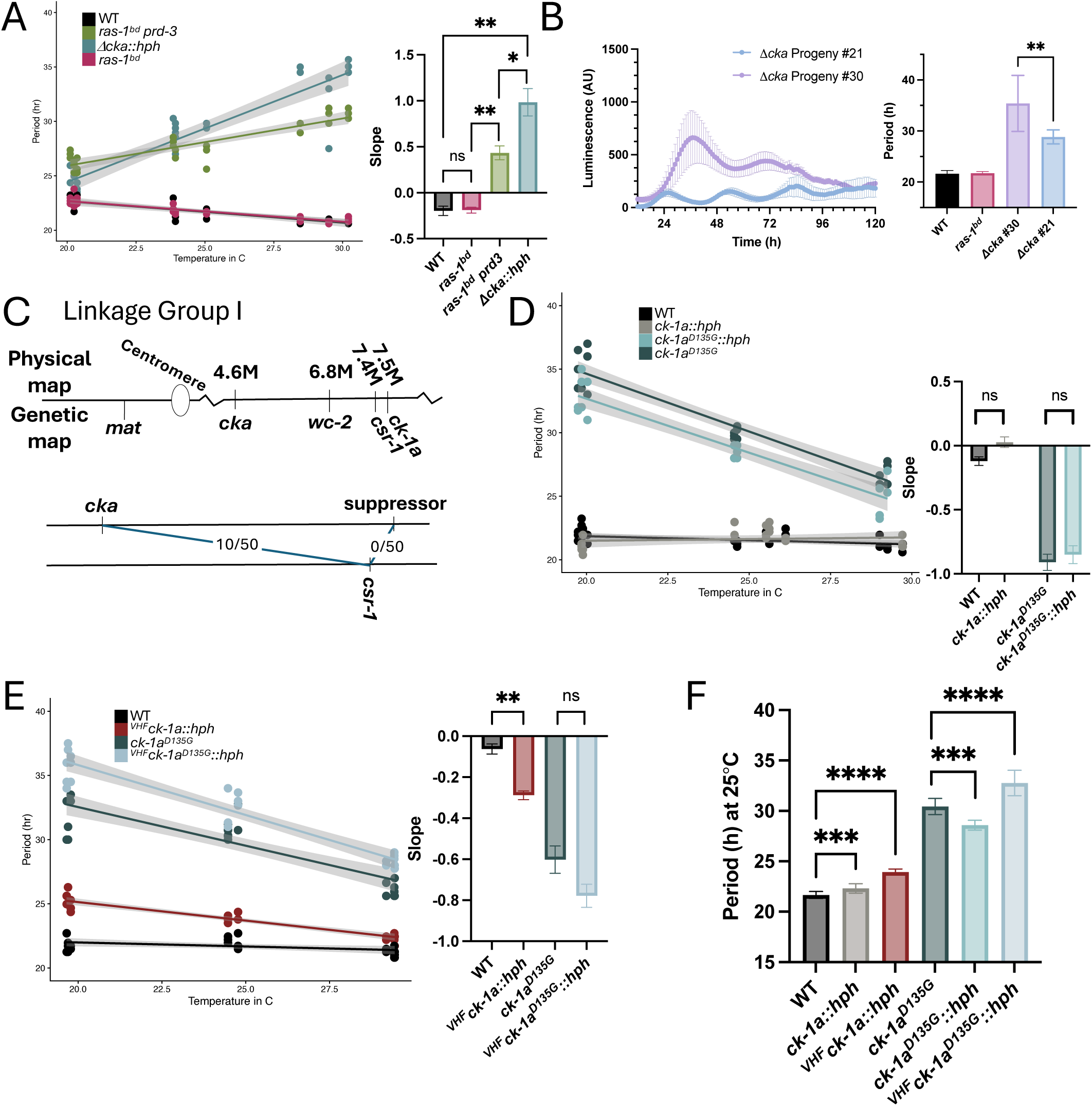
Circadian temperature compensation is regulated by both Casein Kinase I and Casein Kinase II. **(A)** Circadian period length was determined across temperatures for WT, *ras-1^bd^, ras-1^bd^ prd-3* and Δ*cka::hph* using luciferase assays. A linear model was fit to period length, with the grey area showing 95% confidence intervals. The slopes from the linear fit were compared with t-tests. N = 9 at each temperature (3 biological replicates with 3 technical replicates). **(B)** Two siblings from theΔ*cka* backcross to WT are both homokaryotic knock-outs of CKII, but have different period lengths. The trace on the left shows background subtracted luminescence of three technical replicates from one biological replicate at 25°C that was averaged (error bars indicate standard deviation). The bar graph on the right shows the mean period length calculated at 25°C for all replicates +/- standard deviation. Statistical significance was determined by t-test. **(C)** Physical and genetic maps of the right arm of *Neurospora crassa* Linkage Group 1 show close linkage among *ck-1a*, *csr-1* and *wc-2* (above). Recombination frequencies were calculated for cross-overs between *csr-1* and *cka*, and between *csr-1* and *su(Δcka),* later found to be in *ck-1a,* from a backcross of the Δ*cka* strain carrying the suppressor (Δ*cka* #30) to WT. The number of progeny from the backcross with a genotype indicating a crossover event out of the total is listed (below). **(D and E)** Circadian period length was determined across temperatures as in A, for **(D)** WT, *ck-1a::hph*, *ck-1a^D135G^::hph, and ck-1a^D135G^*, and **(E)** WT, ^VHF^*ck-1a::hph*, ^VHF^*ck-1a^D135G^::hph, and ck-1a^D135G^*. **(F)** Period length at 25°C was averaged for strains shown in D and E. Error bars indicate standard deviation. Statistical significance determined by one-way ANOVA.

During the strain construction of *Δcka,* we noticed that two progeny from a backcross to WT, both of which were homokaryotic knock-outs for *cka,* did not have the same period length (**Fig. 1B**), and that one of these siblings was arrhythmic at 20° and 30°C (**Fig. S1B**) while also exhibiting improved growth compared to the other *Δcka* sibling (**Fig. S1C**). Therefore, we hypothesized that a suppressor mutation had arisen in the very sick genetic background of Δ*cka::hph* that happened also to be in a gene affecting the clock. To test this hypothesis, we backcrossed both of these Δ*cka::hph* siblings to WT. As period length was assessed via luciferase reporter, all strains except WT carry the Cbox-luc reporter integrated at the *csr-1* locus. The sibling that was rhythmic at all three temperatures only produced progeny in the backcross that were WT, or Δ*cka::hph* with or without *csr-1::Cbox-luc.* The other sibling that was arrhythmic at 20° and 30°C and had a slightly improved growth phenotype produced progeny that were either WT, [Δ*cka::hph], [*Δ*cka::hph su(Δcka)*] or the [*su(Δcka)*] indicating that the suppressor was not tightly linked to *cka*. While we were able to segregate *su(Δcka)* out to a *cka* WT background, we were not able to separate *su(Δcka)* from *csr::Cbox luc*, indicating tight genetic linkage between *csr-1* and *su(Δcka)* (**Fig. 1C**). Besides *cka*, there are two other core clock genes on Linkage Group 1, where *csr-1* is located: *wc-2* (White Collar 2), a member of the positive arm of the clock, and *ck-1a* (Casein Kinase I, CKI), the kinase that is part of the negative arm of the clock. We sequenced across the entirety of both of these genes using Sanger sequencing and found no changes in *wc-2*, but a point mutation in *ck-1a* that changed aspartic acid at residue 135 to a glycine (**Fig S1D**) in the strains that bore *su(Δcka)*.

We measured the period length of our spontaneous CKI^D135G^ mutation alone, in a *cka* WT background, across temperatures and found it had a very long period and was also extremely under-compensated against temperature with a negative slope in its temperature compensation profile (**Fig. 1D** and **1F**). We confirmed that this mutation was sufficient to cause these circadian phenotypes of a long period and under-compensation by inserting this same mutation into a WT background at the endogenous *ck-1a* locus. We found that our constructed *ck-1a^D135G^::hph* strain had the same circadian phenotypes as the naturally occurring *ck-1a^D135G^* strain (**Fig. 1D**), concluding that *su(Δcka)* is the *ck-1a^D135G^* allele of *ck-1a*. We found that even by only inserting a VHF tag at the N-terminus of WT CKI, the clock is affected with an increase in period length, as well as a modest under-compensation defect (**Fig. 1E – F**). Therefore, the clock appears to be particularly sensitive to changes in Casein Kinase I.

### Temperature compensation regulation is sensitive to changes in CKI and distinct from period length regulation

To further understand the role of CKI in temperature compensation regulation in *Neurospora*, and what may be underlying our *ck-1a^D135G^*mutation, we investigated the impact on temperature compensation of a reduction of *ck-1a* levels, as a reduction in CK1 levels has previously been reported to cause a long period^27^, one of the circadian phenotypes of *ck-1a^D135G^.* Therefore, we replaced the *ck-1a* promoter at its native locus with the *qa-2* promoter, which responds dose dependently to quinic acid^38,39^, in a *ras-1^bd^* (circadian WT^40^) background. We measured the period length of this strain across temperature and a range of quinic acid (QA) concentrations. We found that as we reduced the concentration of QA and had less CKI expressed^27^ (**Fig. S2D**), period lengthened dose dependently (**Fig. 2A, Fig. S2A**). We also found that the clock became under-compensated against temperature at the lowest level of QA tested (statistical significance at 0M QA) (**Fig. 2A**). This indicates that a certain amount of CKI is needed for normal temperature compensation; the clock can handle a reduction of CKI levels but only up to a certain point, at which it can no longer buffer against temperature changes and becomes under-compensated. This is distinct from period length regulation, which was more sensitive to a reduction in CKI levels, with period lengthening with statistical significance starting at 10^-4^M QA. We measured CKI levels by Western Blot at 25°C in *ck-1a^D135G^* and Δ*cka::hph* and found an increase in CKI levels in Δ*cka::hph* compared to WT that approached statistical significance (p=0.0822, one-way ANOVA) (**Fig. 2B**). An increase in CKI levels in Δ*cka* background may explain why a mutation in CKI arose in this background, *ck-1a^D135G^*, that likely reduces CKI function.

**Figure 2.**
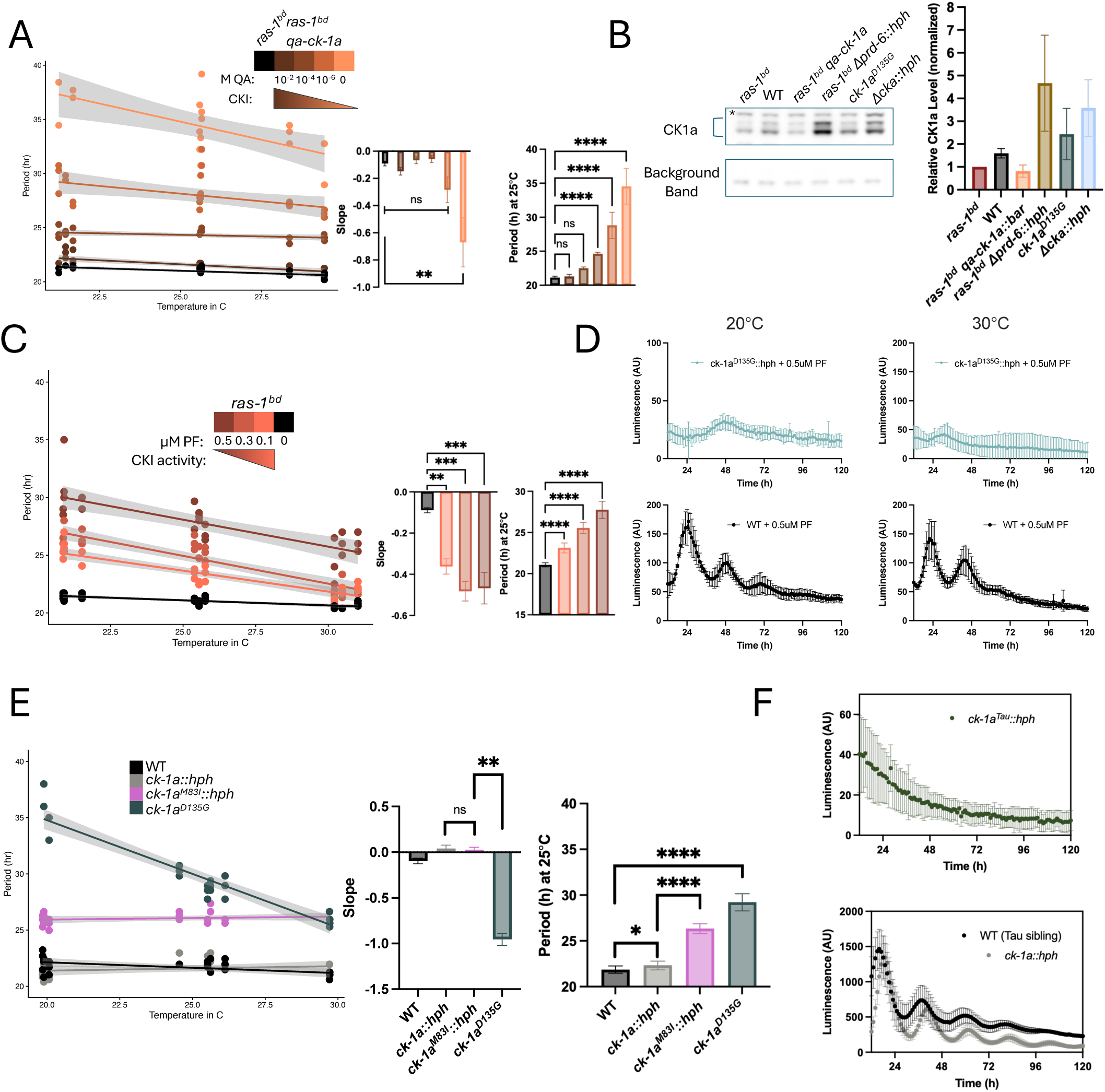
Temperature compensation regulation is sensitive to changes in CKI and distinct from period length regulation. **(A)** The native promoter for Casein Kinase I was substituted with a quinic acid inducible promoter at its native locus, and period length was measured via luciferase assay across temperatures and quinic acid concentrations. A linear model was fit to period length, with the grey area showing 95% confidence intervals. The slopes from the linear fit were compared with a multiple comparison one-way ANOVA (all QA concentrations compared to *ras-1^bd^* control), and the average period length at 25°C of each concentration was compared with a multiple comparison one-way ANOVA. **(B)** CKI levels were assessed by western blot from the indicated strains grown in LCM media (with no quinic acid) at 25°C for 2 days in constant light. A representative example from one replicate is shown (left), and quantification of three replicates is shown (right). CK1 levels were normalized by a background band on each blot, and relative levels were assessed per replicate by setting *ras-1^bd^* to 1. Both isoforms of CKI were included in quantification, indicated by the blue bracket (which does not include the band indicated with *). **(C)** Period length of *ras-1^bd^* was measured across temperatures and concentrations of the CKI inhibitor PF-670462 (“PF”) using a luciferase assay. A linear model was fit to period length, with the grey area showing 95% confidence intervals. The slopes from the linear fit were compared with a multiple comparison one-way ANOVA (all PF concentrations compared to untreated control), and the average period length at 25°C was compared with a multiple comparison one-way ANOVA. **(D)** Representative examples of WT and *ck-1^D135G^::hph* traces from luciferase assay at 20° and 30°C when treated with 0.5uM of PF-670462. An average of three technical replicates from one biological replicate is shown for each genotype and temperature +/- standard deviation. **(E)** Period length of the indicated strains was measured across temperatures using luciferase assays. A linear model was fit to period length, with the grey area showing 95% confidence intervals. The slopes from the linear fit were compared with t-tests. The average period lengths at 25°C are compared with t-tests. The CKI^M83I^ point mutation is at the endogenous locus. **(F)** Representative example of an average of three technical replicates from one biological replicate of WT and *ck-1^R1816^::hph* (Tau) from a luciferase assay +/- standard deviation.

We also found, as others have reported^16,41^, that the clock is sensitive to a reduction in CKI activity in both regulation of period length and temperature compensation. We treated *ras-1^bd^* with the selective CKI inhibitor PF-670462^41^ (“PF”) and found both a dose dependent increase in period length and a dose dependent increase in under-compensation as the concentration of PF increased (and therefore as the activity of CKI decreased) (**Fig. 2C, Fig. S2B**). Unlike a reduction in CKI levels, which became under-compensated only at a very low level of CKI expression, a reduction in CKI activity had an immediate impact on temperature compensation, with under-compensation observed at every concentration of PF that we tested (**Fig. 2C**). Temperature compensation regulation therefore appears to be more sensitive to changes in CKI activity rather than levels. The clock cannot handle the modulation of CKI both by inhibition via PF and the D135G mutation simultaneously, as we observed arrhythmicity when *ck-1a^D135G^::hph* was treated with concentrations of PF that WT was still able to cycle under (**Fig. 2D**). Therefore, the D135G mutation in CKI sensitizes the clock to further reduction in CKI activity. This suggests that while there may be a reduction in CKI activity due to the D135G mutation, there remains *some* amount of CKI activity present in the mutant kinase for the small chemical inhibitor to act on, as we did observe a difference in the clock’s function with and without PF in a *ck-1a^D135G^* background (the clock can still cycle in D135G alone, but cannot cycle with additional PF treatment).

There has been a recent report^16^ of another long period mutant in CKI with an under-compensated clock similar to the CKI^D135G^ strain, CKI^H123Y^. To determine whether a long period phenotype in a CKI mutant always corresponds with dysregulated temperature compensation, we investigated the temperature compensation of the established long period mutant CKI^M83I^, which corresponds to DBT^Long^ in *Drosophila*^42,43^, but whose temperature compensation phenotype has not yet been investigated in *Neurospora*. DBT^Long^ has a long period primarily due to a reduction in kinase activity, and has also been shown to have a small reduction in protein levels compared to WT DBT^42^. The DBT-PER (functionally equivalent to CKI-FRQ) interaction has been determined to be similar between DBT^Long^ and WT DBT^42^. We replaced WT *ck-1a* at the endogenous locus with the *ck-1a* gene followed by hygromycin resistance with no other changes, or with the M83I mutation (**Fig. S2E**), and then measured period length across temperatures using our luciferase clock reporter. While we found the previously reported^44^ long period phenotype in CKI^M83I^, this long period was maintained equally across temperatures, indicating WT temperature compensation (**Fig. 2E, Fig. S2C**). Not only does this indicate that the lack of temperature compensation is more unique in CKI^D135G^ and that not all long period mutants in CKI are necessarily compensation mutants, it also underscores that the regulatory mechanisms that determine period length and those regulating temperature compensation are to some extent distinct.

Given the depth of research into the under-compensation mutant CKIε^Tau^ in the mammalian clock^20,30,34,35^, we generated the corresponding mutation, *ck-1a^R1816^*, in *Neurospora* to see if the phenotypes were conserved. We found that this change in CKI resulted in arrhythmicity (**Fig. 2F**). The strain bearing this mutation was also extremely sick with a very severe growth defect. Therefore, due to the lack of redundancy in *Neurospora* (which only has two isoforms of CKI), we hypothesize that the Tau mutation altered CK1 function to too great a degree to sustain rhythmicity.

### The phosphorylation landscape on FRQ changes with temperature, and in strains lacking normal temperature compensation

Given the drastic changes to temperature compensation we observed following perturbation of either CKII or CKI, two kinases that phosphorylate FRQ, we hypothesized that FRQ phosphorylation may change with temperature, and this may underlie temperature compensation regulation. To test this, we grew V5-6xHis tagged FRQ in *ras-1^bd^*, *ck-1a^D135G^*, and Δ*cka::hph* backgrounds at 20°, 25°, and 30°C in constant light, isolated FRQ for targeted analysis with a two-step pull-down, and then assessed the phospho-occupancy across FRQ using quantitative mass spectrometry (**Fig. 3A**). Three replicates were performed at each temperature, and TMT isobaric labels were used to directly compare across temperature within each genotype. We found that the phosphorylation landscape on FRQ does indeed change with temperature, with higher phospho-occupancy at warmer temperatures in WT (**Fig. 3B**). We found that the phosphorylation landscape on FRQ also changed between genotypes, with a reduction of phosphorylation in Δ*cka::hph*, and higher phosphorylation in *ck-1a^D135G^* at some regions (**Fig. 3B, Fig. S3A**).

**Figure 3.**
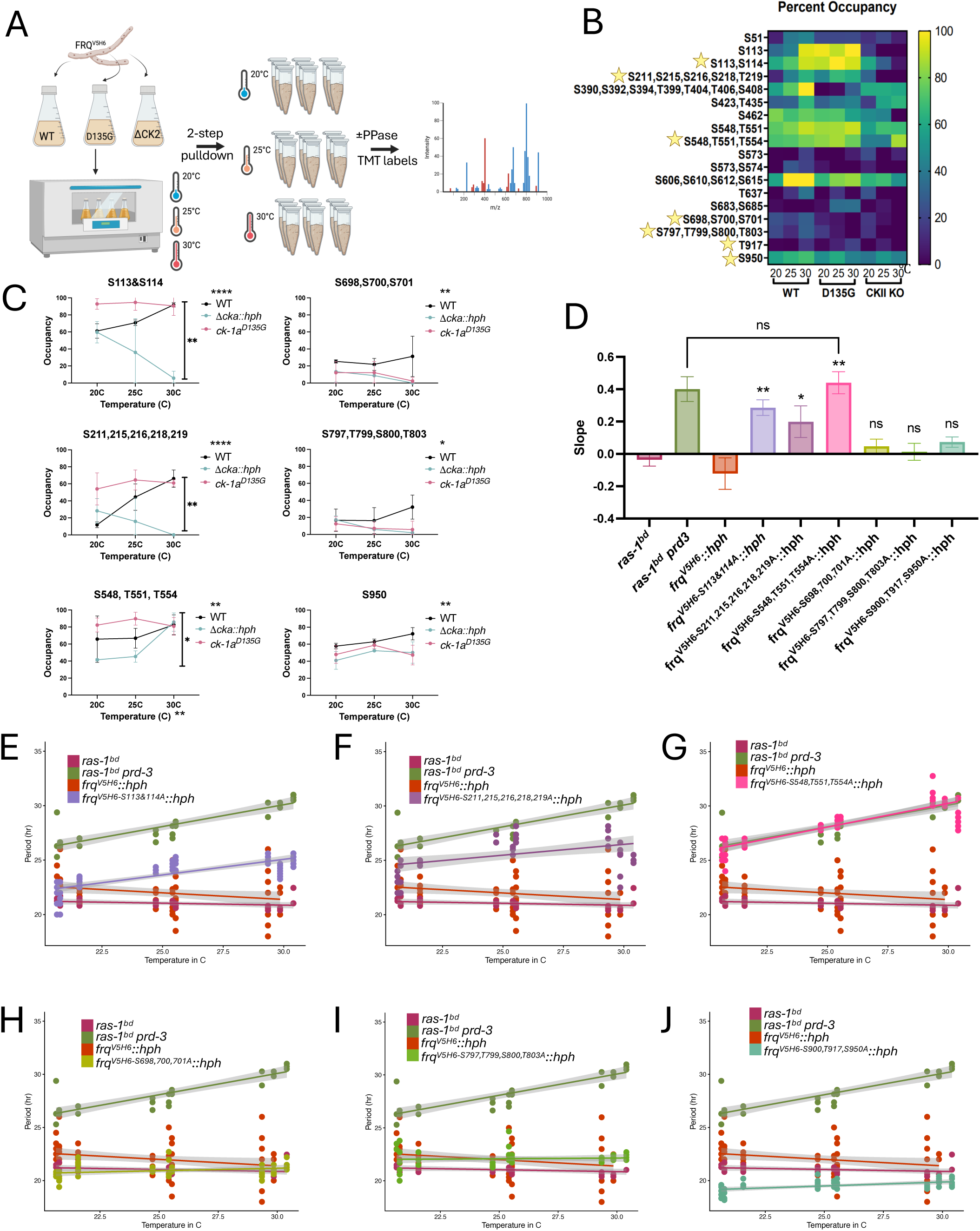
The phosphorylation landscape on FRQ changes with temperature, and in strains lacking normal temperature compensation. **(A)** The experimental set-up to assess quantitative phosphorylation changes on FRQ. A two-step pull-down on FRQ^V5H6^ was performed in a *ras-1^bd^*, *ck-1a^D135G^*, and Δ*cka::hph* background at 20°, 25°, and 30°C (n=3 at each temperature). Phosphorylation status was assessed by treating half of each sample with a phosphatase, and TMT isobaric labels were used to compare across temperatures within each genotype. **(B)** A heat-map of the average phospho-occupancy on FRQ at the indicated regions across three replicates at each temperature in all three genotypes**. (C)** A two-way ANOVA determined the sites at which the phospho-occupancy changed with either temperature (indicated at the x-axis), genotype (indicated at the legend), or the interaction between genotype and temperature (indicated within the plot). Three sites that had a significant interaction between genotype and temperature (left side) and three sites that did not have a significant interaction (right side) are shown. These are the sites that are starred in panel B and were followed up with phosphonull mutations. **(D)** Phosphonull (S/T to A) mutations were constructed at the starred regions, targeted to the endogenous FRQ locus, and then period length was measured across temperature using luciferase assays. A linear model was fit to period length. The slopes from the linear fit are shown +/- standard deviation. Statistical significance was determined with a t-test comparing the phosphonull mutant to *frq^V5H6^::hph* unless otherwise shown**. (E – J)** Temperature compensation profiles from which the slope was determined for panel D are shown for the following changes to *frq^V5H6^::hph:* **(E)** S113CS114A**, (F)** S211, 215, 216, 218, 219A, **(G)** S548, T551, T554A**, (H)** S698, 700, 701A**, (I)** S797, T799, S800, T803A, and **(J)** S900, T917, S950A. Grey area indicates 95% confidence interval.

We performed a two-way ANOVA to determine which phosphosites on FRQ changed with statistical significance across temperature, between genotypes, or had a significant interaction effect of temperature *and* genotype (**Fig. S3B**). The interaction statistic suggests that the temperature dependance of particular phosphosites are different between genotypes. We chose 3 regions (*frq^S113&S114^*, *frq^S211,S215,S216,S218,T219^*, *frq^S548,T551,T554^*) that had a significant interaction effect between genotype and temperature (**Fig. 3C, left**), and 3 regions (*frq^S698,S700,S701^*, *frq^S797,T799,S800,S803^*, *frq^S900,T917,S950^*) at which this interaction was non-significant (**Fig. 3C, right**), and created phospho-null mutations at these sites on FRQ by substituting serine/threonine with alanine. We assessed the period length of these mutants across temperature and found that the sites that had a significant interaction between genotype and temperature had a temperature compensation defect when mutated to alanine (**Fig. 3D –G, Fig. S3CGD**). However, the sites that did not have a significant interaction effect between genotype and temperature maintained WT temperature compensation (despite modest period length changes) when mutated to alanine (**Fig. 3D, H – J; Fig. S3CGE**). The strain with the most severe temperature compensation defect, *frq^V5H6-S548A,T551A,T554A^::hph* had a slope that was not statistically different from the CKII hypomorph, *prd-3* (**Fig 3D, G**).

However, the phospho-occupancy of S548, T551, and T554 was reduced in a *Δcka* background compared to WT only at 20° and 25°C, and had the same occupancy as WT at 30°C, despite the longest period length of *frq^V5H6-S548,T551,T554A^::hph* at 30°C, suggesting that these are likely not CKII sites, and the temperature compensation defect in this strain is due to distinct regulation of temperature compensation from CKII. Overall, the observation that temperature compensation defects were found only in strains that had altered phosphorylation occupancy across temperature (as determined by the interaction statistic) suggests that unique temperature dependent phosphorylation on FRQ may contribute to temperature compensation regulation, and that there are temperature compensation relevant phosphosites on FRQ that are distinct from the sites that just contribute to determining period length.

Some additional limited conclusions are possible from these data. The phospho-occupancy of residues S113CS114 and to a similar extent S211 to S219 on FRQ are consistent with a model where CKII has little impact at 20°C and a much larger impact at 30°C given that the percent occupancy at 20°C is similar in both WT and *Δcka* backgrounds, but at 30°C the occupancy at these sites is greatly reduced in *Δcka*. All three of the phosphonull mutants that had temperature compensation defects show increased occupancy in *ck-1a^D135G^*. If we assume based on previous figures that CK1^D135G^ has reduced activity, then CK1 would be seen as having limited impact at 30°C (as it has a long period at 20°C, but the closest period to WT at 30°C). That these phosphosites are more occupied in CK1^D135G^ suggests that their phosphorylation is repressed by CKI, and yet these cannot be exclusively CKII sites because they remain phosphorylated to some degree in *Δcka*. This suggests the possibility of involvement of additional kinase(s). In the *ck-1a^D135G^* background, we also observe phospho-occupancy maintained evenly across temperature at nearly all sites, compared to the general increase in occupancy at higher temperatures in WT (**Fig 3C, S3B**). This suggests that the temperature compensation defect of *ck-1a^D135G^*could be due to a loss in the temperature dependence of FRQ phosphorylation, in addition to the reduction in activity that leads to long period.

Regardless of which kinase is responsible for the phosphorylation, it is clear that phosphorylation on FRQ plays a role in regulating temperature compensation, as we do not observe only period defects when phosphonull mutations are created: some regions on FRQ (but not all) are important for temperature compensation because temperature compensation is altered when their phosphorylation is prevented by alanine substitutions.

### The impact of CKI inhibition on temperature compensation depends on the amount of CKI and presence of CKII

Given the accumulation of evidence now to suggest that both CKI and CKII have a role in temperature compensation regulation, we next asked what the relationship is between these two kinases in the temperature compensation mechanism: were these kinases acting independently of one another to maintain temperature compensation, or is one dominant over the other? To investigate this, we perturbed both CKI and CKII in the same strain at the same time and then assessed the impact on temperature compensation.

We returned to the small chemical CKI inhibitor, PF-670462 (PF), which causes dose dependent undercompensation in *ras-1^bd^* (WT) (**Fig 2C**). In the over-compensated CKII hypomorph background, *ras-1^bd^ prd-3,* we found again both dose dependent increase in period length and a decrease in slope (towards under-compensation) in its temperature compensation profile with increasing inhibition of CKI via PF treatment (**Fig. 4A, Fig. S4A**), similar to treatment of *ras-1^bd^* with PF. Despite the perturbation to CKII that was already impacting temperature compensation (the *prd-3* mutation), CKI inhibition was still able to alter temperature compensation. This suggests CKI dominance to or co-dominance with CKII in temperature compensation regulation, most preferentially at 20°C where the largest period change is observed with PF treatment, changing period from 24.33h to 31.89h with 0.5μM PF (**Fig. S4E**). At 30°C, CKI inhibition has very little impact on period length in a *prd-3* background, and only modestly lengthens with statistical significance at 0.5μM, changing period from 30.26h to 31.61h (**Fig. S4E**).

**Figure 4.**
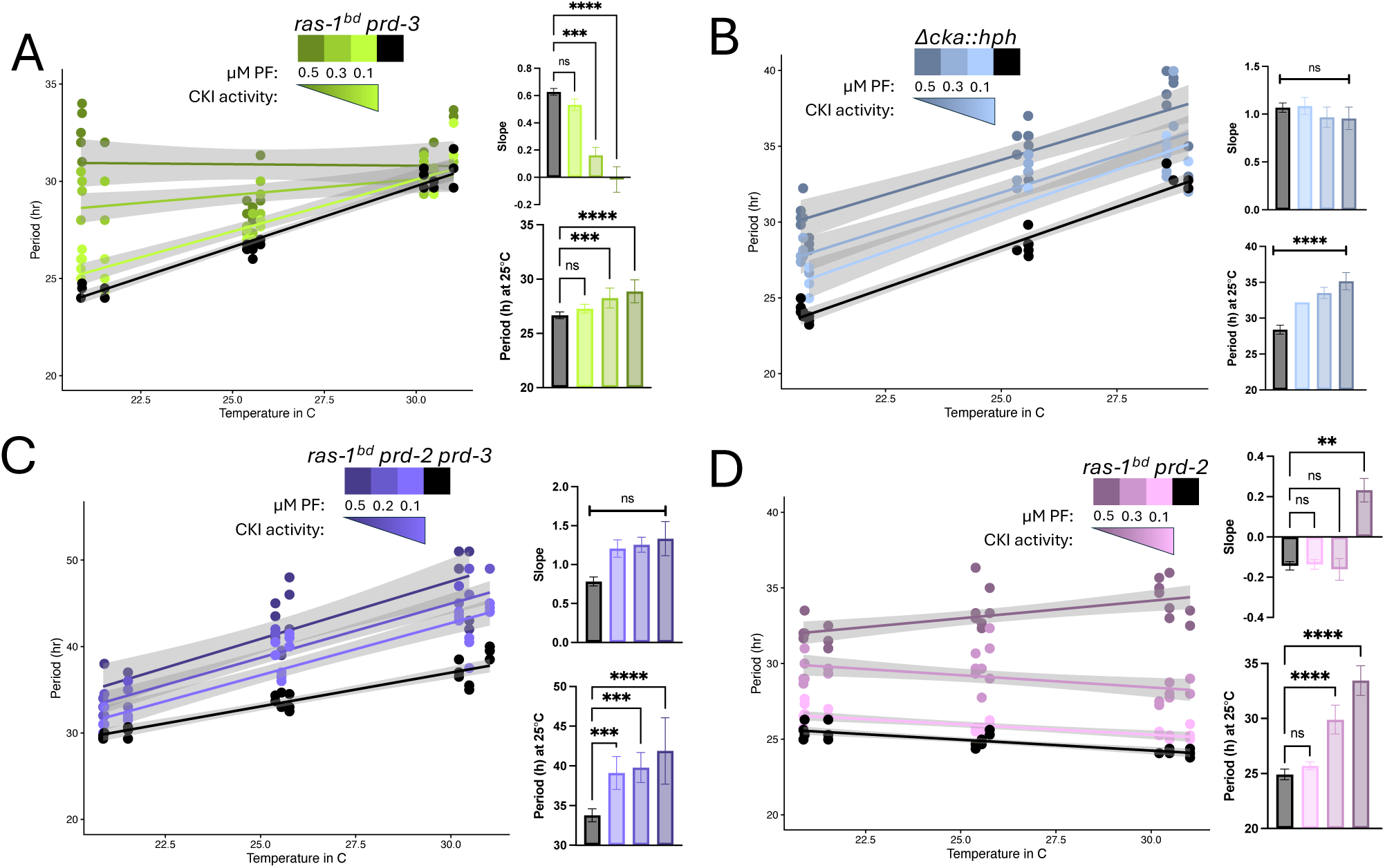
The impact of CKI inhibition on temperature compensation, but not period length, depends on the amount of CKI and presence of CKII. Period length of (A) *ras-1^bd^ prd-3*, **(B)** Δ*cka::hph*, (C) *ras-1^bd^ prd-2 prd-3,* and (D) *ras-1^bd^ prd-2* was measured across temperatures and concentrations of the CKI inhibitor PF-670462 (“PF”) using a luciferase assay. A linear model was fit to period length, with the grey area showing 95% confidence intervals. The slopes from the linear fit were compared with a multiple comparison one-way ANOVA (all PF concentrations compared to control), and the average period length at 25°C was compared with a multiple comparison one-way ANOVA (all PF concentrations compared to control).

However, when we treated the complete knock-out of CKII, Δ*cka::hph*, with this same CKI inhibitor, while we still observed the expected dose dependent increase in period length, CKI inhibition no longer had any impact on temperature compensation (**Fig. 4B, Fig. S4B**). The same over-compensation phenotype of Δ*cka::hph* was maintained at all concentrations of PF, suggesting complete epistasis of Δ*cka::hph* over CKI inhibition for temperature compensation regulation. These data also suggest that CKI may require CKII in order to impact temperature compensation, as when no CKII was present, CKI perturbation no longer effected temperature compensation at all, suggesting CKII is downstream of CKI in the temperature compensation mechanism. CKII protein, even with reduced activity as is the case in *prd-3*, appears to be sufficient for CKI to impact temperature compensation. We note again that this is distinct from period length regulation, as increasing PF concentration still lengthened period dose dependently in theΔ*cka::hph* background, despite having no effect on temperature compensation.

Another known over-compensated strain is the double mutant *prd-2 prd-3*, that has a greater over-compensation defect than *prd-3* alone despite the WT temperature compensation seen in *prd-2* alone (indicating genetic synergy)^45^. We recently identified PRD-2 as an RNA binding protein, knock-out of which results in a reduction in *ck-1a* levels due to a loss in the protection of *ck-1a* mRNA from nonsense-mediated decay^4^^6^. We treated *ras-1^bd^ prd-2 prd-3* with increasing concentrations of PF and observed a dose dependent increase in period length (**Fig. 4C, Fig. S4C**). However, we only observed a modest, non-significant, increase in slope (towards more over-compensation) (**Fig. 4C**), despite the significant <u>decrease</u> in slope (under-compensation) we found when we treated *prd-3* alone with the CKI inhibitor (**Fig. 4B**). Because of this discrepancy, we also treated *ras-1^bd^ prd-2* with increasing concentrations of PF. We again observed the same dose dependent increase in period length with CKI inhibition, but at the highest concentration of CKI inhibition we observed over-compensation in *prd-2,* increasing the slope of its temperature compensation profile (**Fig. 4D, Fig. S4D**). This is more similar to the effect that CKI inhibition had on the temperature compensation profile of *prd-2 prd-3*. Therefore, the impact that CKI inhibition has on *prd-2 prd-3* is likely due to its reduction in CKI levels from, or other impacts of, *prd-2*, rather than its reduction in CKII activity from *prd-3*.

### CKII has a larger role in temperature compensation regulation than CKI

We next turned to genetic manipulation of *ck-1a* in CKII perturbed backgrounds rather than chemical inhibition, to assess the relationship between CKI and CKII in temperature compensation.

In a clock WT, *ras-1^bd^*, background, we found that reduction of CKI levels increases period length, and at very low levels of CKI causes under-compensation (**Fig. 2A**). We used the same quinic acid inducible promoter system driving *ck-1a* in a *ras-1^bd^ prd-3* background and observed an increase in period length with a reduction in CKI levels (**Fig. 5A, Fig. S5A**). However, we saw no impact on temperature compensation as CKI levels decreased, until very low level of CKI expression, where we found extreme over-compensation, the opposite of what we observed in WT (**Fig. 5A**). As over-compensation is a CKII perturbation phenotype, this suggests epistasis of *prd-3* over reduction in CKI levels in temperature compensation regulation.

**Figure 5.**
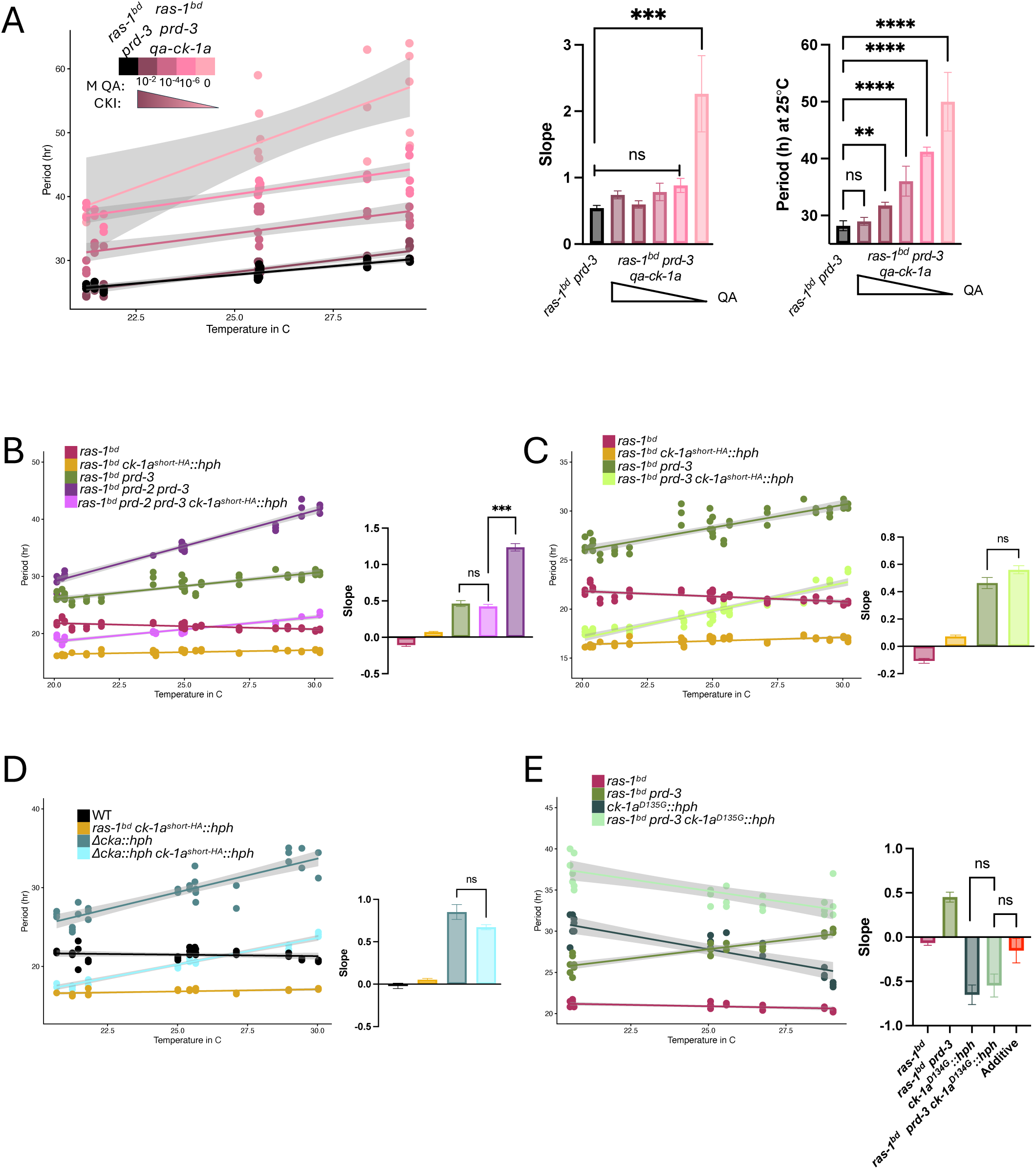
CKII predominantly has a larger role in temperature compensation regulation than CKI. **(A)** Casein kinase I was driven at its native locus with a quinic acid inducible promoter in a *ras-1^bd^ prd-3* strain, and then period length was measured via luciferase assay across temperatures and quinic acid concentrations. A linear model was fit to period length, with the grey area showing 95% confidence intervals. The slopes from the linear fit were compared with a multiple comparison one-way ANOVA (all QA concentrations compared to *ras-1^bd^ prd-3* control), and the average period length at 25°C of each concentration was compared with a multiple comparison one-way ANOVA. **(B – D)** The hyperactive CKI strain, *ras-1^bd^ ck-1a^short^::hph*, was crossed to **(B)** *ras-1^bd^ prd-2 prd-3*, **(C)** *ras-1^bd^ prd-3*, and **(D)** Δ*cka::hph*, and then period length was measured across temperatures using a luciferase assay. A linear model was fit to period length, with the grey area showing 95% confidence intervals. The slopes from the linear fit were compared with t-tests. **(E)** The CKII hypomorph, *ras-1^bd^ prd-3*, was crossed to *ck-1^D135G^::hph* and then period length was measured across temperatures using a luciferase assay. A linear model was fit to period length, with the grey area showing 95% confidence intervals. The slopes from the linear fit were compared with t-tests, and the average period length at 25°C was compared with t-tests. The red bar indicates the slope from a linear fit of the sum of the average period lengthening that *ras-1^bd^ prd-3* contributes at each temperature plus the average period lengthening that *ck-1a^D135G^::hph* contributes at each temperature, including the standard deviations. See Supplemental Figure 5 for the additive temperature compensation profile.

Both PF treatment and the quinic acid promoter system enabled us to reduce overall CKI activity or levels. To look at the impact of changing CKI in the opposite direction, we used the *ck-1a^short^* allele^4^^6^, a hyperactive version of CKI that is missing the auto-inhibitory tail of CKI, and combined this allele with CKII perturbations in the same strain. The *ras-1^bd^ ck-1a^short-HA^::hph* strain alone has a very short period, but maintains this short period evenly across temperatures, indicating normal temperature compensation (**Fig. 5B-D**). We crossed this strain to the *prd-2 prd-3* double mutant and found that the strain that bears all three of these alleles has a much shorter period length than *prd-2 prd-3* alone, and no longer has an extreme over-compensation phenotype (**Fig. 5B, Fig. S5B**).

Instead, this triple mutant returns to a temperature compensation profile similar to *prd-3* alone (**Fig. 5B**, no statistical difference between *prd-3* and *ck-1a^short^ prd-2 prd-3* slopes). The hyperactivity of *ck-1a^short^*appears to fully rescue the change to temperature compensation caused by the reduction in CKI levels from the *prd-2* mutation, leaving only the *prd-3* phenotype in the triple mutant.

We found that both the double clock mutant of *ck-1a^short^* and *prd-3* (**Fig. 5C, Fig. S5C**), as well as the double clock mutant of *ck-1a^short^* and Δ*cka::hph* (**Fig. 5D, Fig. S5C**) have a shortened period length compared to *prd-3* or Δ*cka::hph* alone, but maintain the same over-compensation phenotypes of the single mutants *prd-3* and Δ*cka::hph*, respectively. This indicates epistasis of *prd-3* and Δ*cka::hph* over *ck-1a^short^* for temperature compensation regulation, and therefore dominance of CKII over CKI for temperature compensation. Importantly, the over-compensation phenotype observed in the double mutants of *ras-1^bd^ prd-3 ck-1a^short-HA^::hph* and Δ*cka::hph ck-1a^short-HA^::hph* is **not** simply due to additive period length contributions of *prd-3* or Δ*cka::hph* plus the contribution of *ck-1a^short-HA^::hph,* which would indicate these alleles are independently regulating temperature compensation; this is not the case here (**Fig. S5D**).

We assessed the genetic relationship between *ck-1a^D135G^*and *prd-3* by combining these mutations in the same strain and measuring period length across temperatures. We found the double mutant bearing both *ck-1a^D135G^* and *prd-3* had a period length that was the sum of the period lengthening caused by *ck-1a^D135G^* and the period lengthening caused by *prd-3* at both 25°C and 30°C (**Fig. S5G**), indicating regulation via separate pathways at these temperatures. However, at 20°C, the period length was longer than what would be expected from additive (**Fig. S5G**), making the double mutant under-compensated (**Fig. 5E, Fig. S5EGF**). The under-compensation found in the double mutant was not significantly different than the temperature compensation profile of what is expected from just additive period lengthening at each temperature (**Fig. 5E**). However, the slope of the double mutant was also not significantly different from the slope of *ck-1a^D135G^::hph* alone (**Fig 5E**). This again suggests a greater role for CKI than CKII at 20°C, as we observed under-compensation in the double mutant (a *ck-1a^D135G^*like phenotype).

## Discussion

Both CKI and CKII contribute to temperature compensation in *Neurospora*, and our data have further defined their roles. The complete knock-out of CKII is even more overcompensated than its corresponding hypomorph, *prd-3*, and to our surprise, a suppressor mutation that arose in this background, *ck-1a^D135G^*, not only impacts the clock by lengthening period, but impacts temperature compensation as well, with an effect opposite to Δ*cka*. A model begins to emerge in which CKI has a larger role in period length maintenance at 20°C, and CKII has a larger role in period length maintenance at 30°C. The phosphorylation dynamics within the clockwork change across temperatures, creating a scenario where the relative importance of each kinase changes dramatically with temperature (**Fig. 6**). This may explain the changes in the FRQ phosphorylation landscape at different temperatures that we observed in our quantitative phosphoproteomics. When CKII is disrupted, period lengthens away from WT more at 30°C than 20°C, suggesting the clock relies on CKII more at 30°C. When CKI is disrupted, from either mutations or a reduction in levels or activity, the period lengthens away from WT much more at 20°C than 30°C, suggesting the clock relies on CKI less at 30°C, and more at 20°C. Biochemical investigations suggest that CKI is inherently temperature compensated^15,24^, unlike most kinases, while CKII does increase its activity with temperature^27^. The increase in FRQ protein at higher temperatures^47^ may require greater kinase activity to phosphorylate a higher number of FRQ proteins. Because of this, CKI (which does not increase its activity with temperature) may require the help of another kinase, such as CKII (which is more active at higher temperature), in order to phosphorylate all of the available FRQ protein at higher temperatures, but would not require as much help at lower temperature where there is less FRQ protein present.

**Figure 6.**
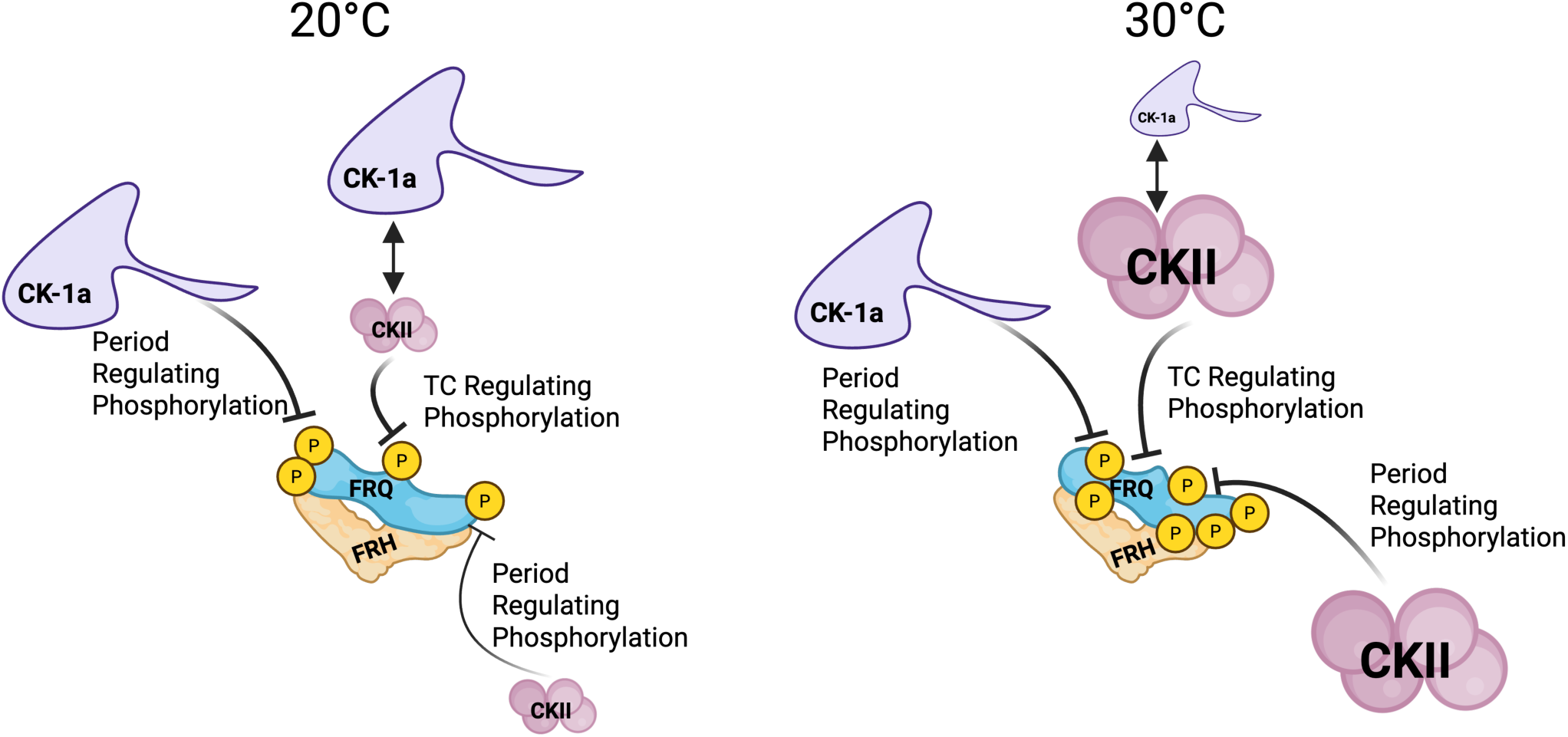
A working model for temperature compensation regulation mediated by Casein Kinase I and II. CK1a perturbations cause a greater change in period length at cold temperatures than warm temperatures and thus have a larger role in regulating temperature compensation (TC) at cold temperatures than at warm temperatures, likely through FRQ phosphorylation. As CK1a activity does not increase appreciably with temperature, it would perform period-regulating phosphorylation on FRQ at similar levels at cold and warm temperatures. Casein Kinase II (CKII) has a larger role in regulating temperature compensation at warm temperatures, as CKII perturbations cause a great change in period length at warm temperatures, and only very small period changes at cold temperatures. CKII activity exhibits normal temperature dependence and therefore increases its activity at warm temperatures, phosphorylating more at these temperatures. CK1a may act through CKII to impact temperature compensation by CKII mediated increase in the affinity of CKI for FRQ at high temperature. These kinases regulate temperature compensation via phosphorylation at specific regions/phosphosites on FRQ that impact temperature compensation. This change in the dominant kinase phosphorylating FRQ as temperature changes is accompanied by changes in the phosphorylation landscape on FRQ as temperature changes (i.e., which phosphosites are occupied), as well as increasing the quantity of phosphorylation as temperature increases: higher phospho-occupancy is found on FRQ at higher temperatures. This is consistent with more kinase activity on FRQ at warm temperatures, driven by consistent CK1a activity across temperatures but increased CKII activity at warm temperatures, resulting in net higher kinase activity at warm temperatures and consequently more phosphorylation on FRQ at higher temperatures. This increase in phosphorylation on FRQ may work to counteract the increased protein level of FRQ at higher temperature.

Our treatment of *prd-3* with the CKI inhibitor, PF-670462, is consistent with CKI dominance in period regulation at cold temperatures more so than at high temperatures and CKII dominance instead at high temperatures, as we observed 7 hours of period lengthening at the highest concentration of PF at 20°C, but just over an hour of period lengthening with the same concentration at 30°C. Zooming in on just the 30°C data in this instance, *prd-3* is epistatic to CKI inhibition at all concentrations except for 0.5μM PF where we observe the modest one hour increase in period length (**Fig. S4E**). Surprisingly, this temperature dependence of the degree of period lengthening with CKI inhibition disappears when CKII is not physically present in the CKII knock-out, suggesting CKI acts *through* CKII for temperature compensation (but not for general period regulation, meaning CKI can still phosphorylate FRQ at some regions without CKII), as we found no change in temperature compensation in Δ*cka::hph* upon PF treatment. CKI was still able to influence temperature compensation when CKII protein was present, but had reduced activity in a *prd-3* background. It is possible that CKI may not need the catalytic function of CKII in order to influence temperature compensation. Instead, CKI may use a non-catalytic role^48^ of CKII to mediate temperature compensation, such as using CKII as a physical scaffold, or CKII regulating CKI allosterically.

Many phosphonull mutations on FRQ have been reported^49–51^ that alter period length, and a small subset of these also change the temperature compensation profile of the clock. Importantly however, not every mutation leading to period changes is accompanied by a change in temperature compensation. Consequently, there must be phosphosites or regions on FRQ that are specific to temperature compensation regulation rather than just regulating period (**Fig. 6**). We found that the sites on FRQ whose phospho-occupancy in response to temperature differed in strains with temperature compensation defects were temperature compensation specific sites. When alanine was substituted at these positions, the rate of the clock was no longer maintained within a physiological temperature range. This change in FRQ phosphorylation with temperature may be due to the change in dynamics between CKI and CKII with temperature, with one kinase dominant, or needed, at cold temperature, and the other kinase dominant at warm temperature. The phosphorylation occupancy data also suggested that CK1 may play a repressive role at compensation-relevant sites as the reduction of CKI activity in CKI^D135G^ was accompanied by increased occupancy at these sites. These data are also compatible with the action of CKI and CKII on an additional unknown kinase(s). We also observed a loss in the temperature dependence of FRQ phosphorylation in *ck-1a^D135G^,* which has a severe temperature compensation defect, further underscoring that dynamic, temperature dependent, phosphorylation of FRQ contributes to temperature compensation maintenance.

We noted that temperature compensation was not impacted in the exact same way from a reduction in CKI activity as a reduction in CKI levels (**Fig. 2**). Specifically, a reduction in CKI activity had a much larger and immediate impact on temperature compensation. We speculate that this could be because there is a specific, clock relevant, pool of CKI that is important for temperature compensation. If we reduce global CKI levels it may take a larger reduction to reach the limiting factor on CKI levels for circadian function. However, if we reduce CKI *activity* globally, even the clock relevant pool that is being used in the clock at that very moment is affected by the inhibitor, so we see under-compensation more dramatically and more rapidly. This is consistent with recent data^52,53^ that suggests that FRQ forms condensates that recruit other clock components, including CKI, which could distinguish this clock relevant CKI pool from the rest of available CKI.

A current model for temperature compensation and period length determination in *Neurospora* suggests that the FRQ-CKI interaction underlies both of these mechanisms^16,29^ and their regulation therefore go hand in hand. However, as noted above, we found that the relationship between CKI levels and either period length or compensation is not rigidly fixed: WT temperature compensation persists far longer than WT period length when CKI levels are reduced. According to that model, this reduction in CKI levels is weakening the FRQ-CKI interaction to increase period length, but somehow this weaker-than-WT FRQ-CKI interaction is still maintained equally across temperature, until CKI levels reach a critically low threshold. We find that the CKI^M83I^ mutant, equivalent to DBT^Long^ in *Drosophila,* also contradicts the FRQ-CKI interaction model, as it showed a WT temperature response despite a long period. As DBT^Long^ has been shown to have reduced kinase activity^42^, which we also show *does* alter temperature compensation, there may be something compensating for this reduction in activity to prevent further period changes across temperature. DBT^Long^ is able to maintain an interaction with PER similar to WT DBT^42^. If this interaction is maintained equally across temperature, it may explain the lack of a temperature compensation defect, as the FRQ-CKI (equivalent to PER-DBT) model predicts that WT FRQ-CKI interaction should maintain WT TC^16^. However, while the temperature compensation phenotype of CKI^M83I^ is consistent with the FRQ-CKI interaction model, the long period would not be predicted by a model that FRQ-CKI interaction also determines period length^29^, because DBT^Long^ has a long period, but normal PER-DBT interaction. This interaction was only tested in *Drosophila*, however, so it is possible the FRQ-CKI interaction might be different in *Neurospora* for this mutant. The FRQ-CKI interaction model for temperature compensation implies that TC is regulated by modulating the affinity of CKI for FRQ, which would alter the phosphorylation of all of FRQ in general and all phosphosites on FRQ are treated alike. However, we find that there are distinct phosphosites on FRQ that impact period alone, and others that impact both period length and temperature compensation, contradicting this implication.

It has been shown that a CKII hypomorph, *ckb^RIP^*, reduces the FRQ-CKI interaction^16^. A reduction in CKII activity leading to a reduction in the affinity of CKI for FRQ suggests that under normal, WT conditions, CKII is increasing the FRQ-CKI interaction, and does so even more at high temperature. This could be done either catalytically, such as by CKII dependent FRQ phosphorylation altering FRQ conformation^54^ that favors CKI binding, or non-catalytically as discussed above. Some of our data is consistent with this, as a weaker FRQ-CKI interaction at high temperatures would result in a longer period at higher temperatures (i.e. overcompensation as seen in *prd-3* and Δ*cka::hph*). This is also consistent with the lack of temperature dependence in period lengthening conferred by CKI inhibition in Δ*cka::hph* when CKII is not present: CKI disruption evenly increases period length across temperature, as CKII is not increasing the affinity of CKI for FRQ at higher temperature, leaving only the TC defect from the CKII knock-out.

We also find that temperature compensation is never altered without also altering period length: we have not observed a WT period length at 25°C (the temperature at which period is measured when not assessing TC) when temperature compensation is disrupted. While absence of evidence is not evidence of absence, this does suggest that the TC regulating phosphorylation on FRQ is likely more robust than the period regulating phosphorylation on FRQ. That is to say, a mutation is far more likely to disrupt period length than TC, and if a mutation disrupts the clock enough to have an impact on temperature compensation regulation, it has already surpassed what is required to disrupt period length regulation. Mechanistically, this could be achieved through a difference in the affinity of CKI for domains on FRQ that regulate period length to those that regulate TC. Indeed, the affinity of CKI is far greater for primed sites than unprimed sites on FRQ, although it is capable of phosphorylating both^24^. If CKI has a weaker affinity for period regulating phosphorylations on FRQ and a greater affinity for temperature compensation regulating phosphorylations on FRQ, period length will necessarily be altered if CKI is disrupted to a great enough degree that temperature compensation is impacted. This is consistent with our finding that there are phoshphosites on FRQ that when mutated to alanine disrupt only period length, while other phosphosites disrupt both period length and temperature compensation (**Fig. 3**), as well as our finding that temperature compensation persists far longer than period length when CKI levels are reduced (**Fig. 2**).

Our data also suggest that the synergistic temperature compensation phenotype observed the *prd-2 prd-3* double mutant^45^ is primarily due to the alteration to CKI levels that is caused by the *prd-2* mutation. We were able to restore the temperature compensation phenotype of the double mutant back to that of *prd-3* alone by creating a triple mutant of *prd-2 prd-3 ck-1a^short^*, which increases the activity of CKI and could compensate for a reduction in CKI levels. Additionally, we forced a reduction in CKI levels in a *prd-3* background using a quinic acid inducible promoter driving *ck-1a*, and observed extreme over-compensation at very low levels of *ck-1a* expression in a *prd-3* background, reminiscent of the *prd-2 prd-3* compensation phenotype^45^. Interestingly, CKI inhibition of *prd-2* increased period length dose dependently, despite low, but present, CKI levels in this strain^46^. The existent CKI in *prd-2* is still enough to keep the clock running (as *prd-2* is not arrhythmic) and still enough that a reduction in activity still has an impact on period. This low CKI level, or other alterations caused by the loss of the RNA-binding protein PRD-2, did change the impact that CKI inhibition had on temperature compensation, however, because the increasing under-compensation with CKI inhibition that occurred in WT (*ras-1^bd^*) was not observed in a *prd-2* background.

In the mammalian clock, phosphorylation dynamics on PER also underlies temperature compensation, mediated by a phospho-switch on PER^30,32^ that is carried out by CKIδ/ε^55^. A similar phospho-switch has been described on *Drosophila* PER that may also underlie temperature compensation in that system^56^, likely from DBT (the CKI homolog) mediated phosphorylation^56–58^. These models rely on a change in PER stability as the outcome of changes in PER phosphorylation, however, which is not the salient outcome of phosphorylation of FRQ in *Neurospora*^17^, despite the presence of degrons on FRQ^59^. FRQ additionally does not have a known stabilization domain, another requirement of the phospho-switch model^60^. Therefore, there may be a modest deviation in functional conservation in the temperature compensation mechanism between the TTFL based clocks (mammals, insects, and fungi), or, the link between these mechanisms remains uncovered. The functional consequence of phosphorylation on PER for temperature compensation (rather than period length determination) may not be changes in PER stability, as is the case in *Neurospora*^17^, since recent data has questioned the degradation based model in mammals^18^. Alternatively, a phospho-switch may instead also exist in *Neurospora*, causing a change in clock regulation directly from changes in phosphorylation rather than changes in stability.

All current TTFL models for temperature compensation have revealed a role for CKI^16,30,56^. CKII is known to affect the clock in mammals, with CKII knock-down causing a long period^61^, but its role in temperature compensation has not been investigated. In *Drosophila*, reduction in CKII also increases period length^62^, similar to the phenotype in both the mammalian clock and the *Neurospora* clock, but its role in temperature compensation is also unknown. Given the conservation in the period length phenotype with CKII perturbation across the TTFL-based clocks, it is likely that conservation in its role in temperature compensation regulation exists as well. Even the plant clock, which involves many interlocking negative feedback loops and is evolutionarily distinct from the *Neurospora* clock, relies on CKII in part to regulate temperature compensation^63^. Therefore, although the mechanism by which these players maintain temperature compensation may deviate between systems, there remains conservation in the players themselves.

## Methods

### *Neurospora* strains

See Supplemental Information for a full list of the *Neurospora* strains used in this study. Homokaryons of Δ*cka::hph* containing the luciferase reporter were generated by crossing the Δ*cka::hph* heterokaryon previously reported^27^ to WT *csr-1::frq_cbox_-luc::bar*. Due to extremely slow growth, the resulting homokaryons took many more days than typical to generate enough growth to be seen by the naked eye, which is likely why they were not generated during previous study^27,64^. The *ck-1a::hph, ck-1a^D135G^::hph*, *ck-1a^M83I^::hph, ck-1a^R1816^::hph, ^VHF^ck-1a::hph,^VHF^ck-1a^D135G^::hph, frq^V5H6^*::hph, frq*^V5H6-S113&S114A^::hph, frq^V5H6-S211,215,216,218,219A^::hph, frq^V5H6-S548,T551,T554A^*::hph, frq*^V5H6-S698,700,701A^::hph, frq^V5H6-S797,T799,S800,T803A^::hph, and frq^V5H6-S900,T917,S950A^::hph* strains were constructed by transforming DNA with 1kb flanking the target into Δ*mus-51::bar* strains to generate heterokaryons. Heterokaryons were crossed to a WT strain containing a circadian luciferase reporter where the “clock box” *frq* promoter drives expression of codon-optimized firefly luciferase^17,37^, inserted at the *csr-1* locus, *csr-1::frq_cbox_-luc::bar*, to generate homokaryons. Presence of the luciferase reporter was determined by cyclosporin resistance and luminescence signal, point mutants were determined by sanger sequencing of either *ck-1a* or *frq* and by hygromycin resistance, and tags were confirmed by western blot.

Double mutants were generated by sexual crosses. Homokaryons carrying both mutations were confirmed by sanger sequencing for *ck-1a^D135G^::hph*, PCR genotyping for Δ*cka::hph*, PCR genotyping or sanger sequencing for *prd-3*, sanger sequencing or presence of HA tag via western blot for *ck-1a^HA-short^::hph*^46^, and CKI expression response to QA for *qa-ck-1a::hph*^27,46^ via CKI western blot.

### PCR Genotyping and Sanger Sequencing

*Neurospora* genomic DNA was extracted using Mouse Tail buffer (Allele Biotechnology # ABP-PP-MT01500). For PCR genotyping, Go Taq Green (Promega M712C) polymerase was used, and for amplifying *wc-2, ck-1a* or *frq* for sanger sequencing, Phusion Flash High Fidelity (ThermoFisher #F548S) polymerase was used. The following primers were used for either genotyping, gene amplification, or for sanger sequencing:

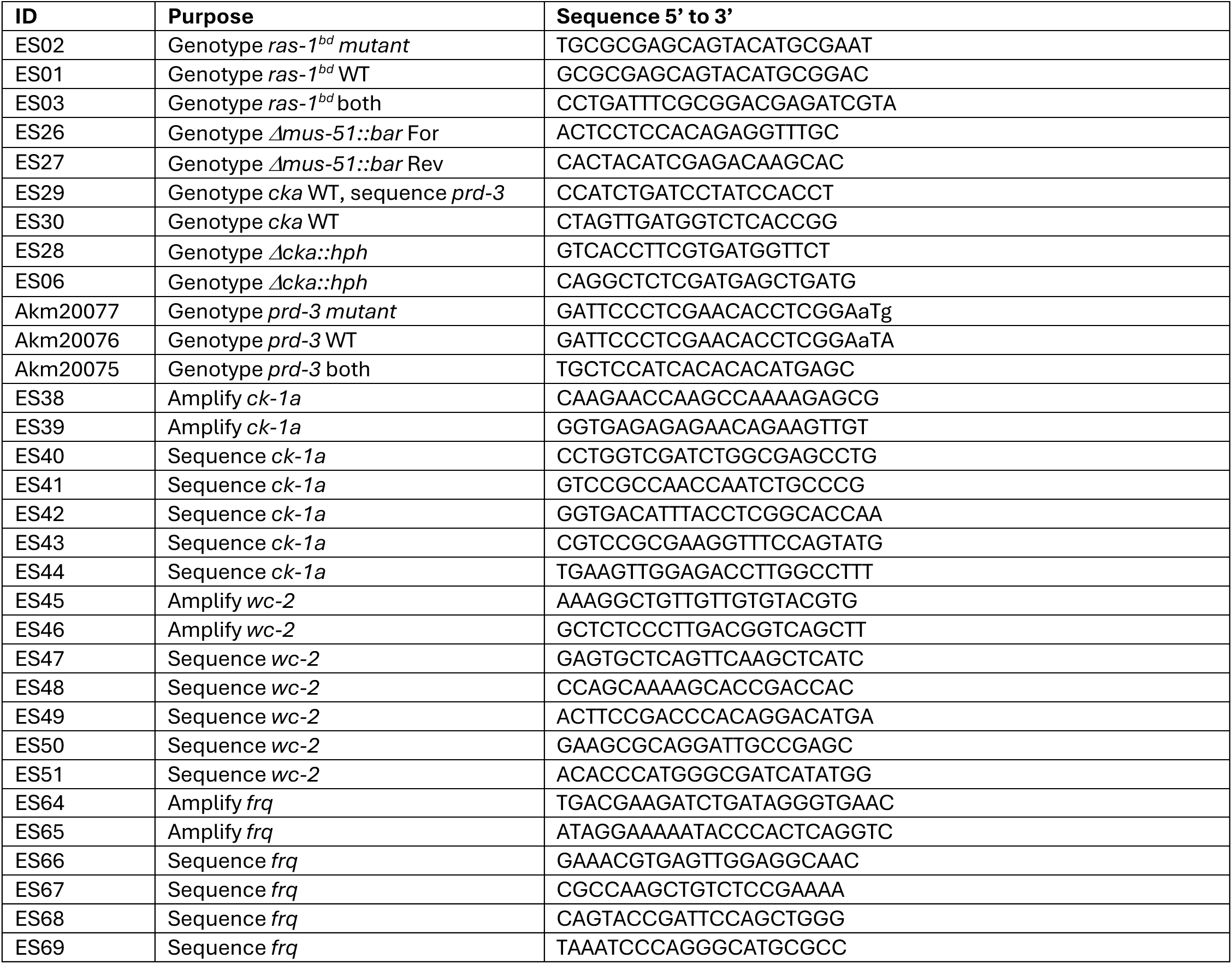

### Circadian Luciferase Assay To Determine Period Length Across Temperature

*Neurospora* conidial suspensions in water were inoculated into 96 well plates consisting of black tubes to prevent light leakage between wells, containing 0.03% glucose 0.05% arginine media with luciferin. For camera runs with CKI inhibitor PF-670462 (Sigma #SML0795) or quinic acid (Sigma #138622), base media was made for the entire plate, and aliquots of this base media was then used to make the varying concentrations of PF or QA media. Three technical replicates of each strain were inoculated into each plate. Plates were entrained at 25°C in a 12-hour light dark cycle for 2 days in a Percival incubator and then transferred into constant darkness at either 20°, 25°, or 30°C in a Percival incubator for the camera run. A HOBO temperature and light monitor (Onset # MX2202) was used to determine the exact temperature of entrainment and camera runs, and the average temperature for the duration of the run was plotted on the x-axis of all temperature compensation profiles. A Pixis 1024B CCD camera was used to collect bioluminescent signal from the luciferase reporter with an exposure time of 15 minutes, repeated once per hour over the course of the run.

Period length was determined by running background-corrected luminescent traces (determined by an ImageJ macro^17^) through an R program that calculates spectral density, as previously described^46^. Several strains reported here have clocks that produced luciferase traces that were less robust than WT so the R program could not accurately determine period, or had a period length that was longer than the R program would accurately call period. For these strains that period length could not be determined by the spectral density R program, period length was determined instead by manually finding the hour at each peak of luminescence, and averaging the time between peaks (peak to peak period call). If the manually determined period call matched the period call determined by the R program, the period call from the R program was used, as it calculates period length using every point, rather than just the peaks. A separate R code, “peakyFinders”, was written to assist in manually determining the peak of bioluminescence that: smoothed the data with the gaussianSmooth function, used the findpeaks function to determine the most likely peak, and then locally around this predicted peak determined the un-smoothed, background subtracted, value with the highest luminescence, and reported the hour this was recorded. Every trace that was run through this peak-to-peak R program was manually confirmed afterwards that the correct peak was called.

Temperature compensation was evaluated by plotting actual temperature of each run (from HOBO monitor) vs the period length determined at that temperature for each technical replicate. A linear regression was fit to this data in Prism GraphPad, and the slope from the linear regression was used to compare temperature compensation profiles across strains. Temperature compensation plots shown in figures were generated in R using ggplot with a linear model fit to the data, and 95% confidence interval around this linear fit is shown in grey.

### Quantitative Phosphoproteomics on FRQ

*ras-1^bd^ frq^V5H6^::bar* was crossed to *ck-1a^D135G^* and Δ*cka::hph* to generate tagged FRQ in these genetic backgrounds. After strain validation, two-step pulldowns were performed on FRQ from these strains by first using nickel resin to pull on the 6xHis tag, and then a Mouse anti-V5 antibody (BioRad MCA1360) on Protein G Dynabeads (Thermo 1003D) was used to pulldown on the V5 tag from the eluate of the nickel pulldown. Tissue was grown in LL at 20°, 25°, and 30°C in 1000mL LCM per replicate, and protein was extracted in Protein Extraction buffer (50 mM HEPES (pH 7.4), 137 mM NaCl, 10% glycerol v/v, 0.4% NP-40 v/v, and cOmplete Protease Inhibitor Tablet (Roche #11 836 170 001). Before proteomics analysis, samples were reduced and alkylated, and pulldown success was confirmed by Western Blot with Rabbit anti-FRQ.

Proteins were enriched with SP3 and digested overnight at 37°C in 133 mM EPPS, pH 8.5, with trypsin (Promega). Samples were split in half and dephosphorylated with CIP (NEB) or not for 2 hours at 37 °C, then digested with trypsin again for 1 hr at 37 °C. Peptides were labeled with Tandem-Mass-Tag (proTMT) reagent (ThermoFisher Scientific). Once labeling efficiency was confirmed to be at least 95%, each reaction was quenched by the addition of hydroxylamine to a final concentration of 0.25% for 10 minutes, mixed, acidified with trifluoroacetic acid (TFA) to a pH of about 2, and desalted over an Oasis HLB plate (Waters). The desalted multiplex was dried by vacuum centrifugation and separated by offline Pentafluorophenyl (PFP)-based reversed-phase HPLC fractionation as previously described^65^. TMT-labeled peptides were analyzed on an Orbitrap Lumos mass spectrometer (ThermoScientific) equipped with Vanquish Neo liquid chromatography system^66^ (ThermoScientific).

The raw data files were searched using COMET with a static mass of 229.162932 Da on peptide N-termini and lysines and 57.02146 Da on cysteines and a variable mass of 15.99491 Da on methionines and 79.96633 Da on serines, threonines and tyrosine against the target-decoy version of the human proteome sequence database (UniProt; downloaded 2/2013, 40,482 entries of forward and reverse protein sequences), maximum of three missed cleavages allowed, precursor ion mass tolerance 1 Da, fragment ion mass tolerance ±8 ppm, and filtered to a <1% FDR at the peptide level. Occupancy was calculated based on peptide intensity in the CIP-treated sample divided by peptide intensity in the control-treated sample and divided by peptide intensity in the CIP-treated sample. P-values were calculated using a two-tailed Student’s t test, paired values.

### Protein Extraction and Western Blot

For Western Blot and two-step pulldown for quantitative proteomics, *Neurospora* protein was extracted as previously described^67^. Briefly, *Neurospora* liquid culture was vacuum filtered, semi-dry tissue was ground in liquid nitrogen, and protein was extracted on ice using PEB (50 mM HEPES (pH 7.4), 137 mM NaCl, 10% glycerol v/v, 0.4% NP-40 v/v, and cOmplete Protease Inhibitor Tablet (Roche #11 836 170 001). Protein amount was quantified using a Bradford Assay (Bio-Rad #500–0006). For the blot in Fig 2B, conidia were inoculated into 3mL of LCM in 35mm dishes overnight. Mycelia mats were then transferred to 50mL of LCM and rotated at 125rpm for two days in LL at 25°C, and then tissue was harvested. 30ug of protein was loaded into 4-12% Bis-Tris gels (Thermo #NP0321BOX) and then transferred to a PVDF membrane. CK1 levels were assessed with a Rabbit Anti-CK1a antibody, gifted from the Brunner Lab.

## Acknowledgments

We thank Dunlap-Loros lab members for helpful discussions. We thank Adrienne Mehalow for *prd-3* genotyping primers. We thank David Ritz for development of peak-peak period call R code, peakyFinders. We thank Axel Diernfellner and Michael Brunner (University of Heidelberg) for generously sharing the *Neurospora* CK1a antibody. This work was supported by NHLBI F31HL165774 to ELS, NIGMS F32GM128252 and R35GM157067 to CMK, NIGMS R35119455 to ANK, and NIGMS R35GM118021 to JCD. The authors declare no conflicts of interest. Figures 3A and 6 were created in BioRender. Stevenson, E. (2026) https://BioRender.com/nzqvt9g and https://BioRender.com/tu332jq.

