## Supplemental Figures for "The Case for Kinases: A Phosphorylation Driven Model for Circadian Temperature Compensation"

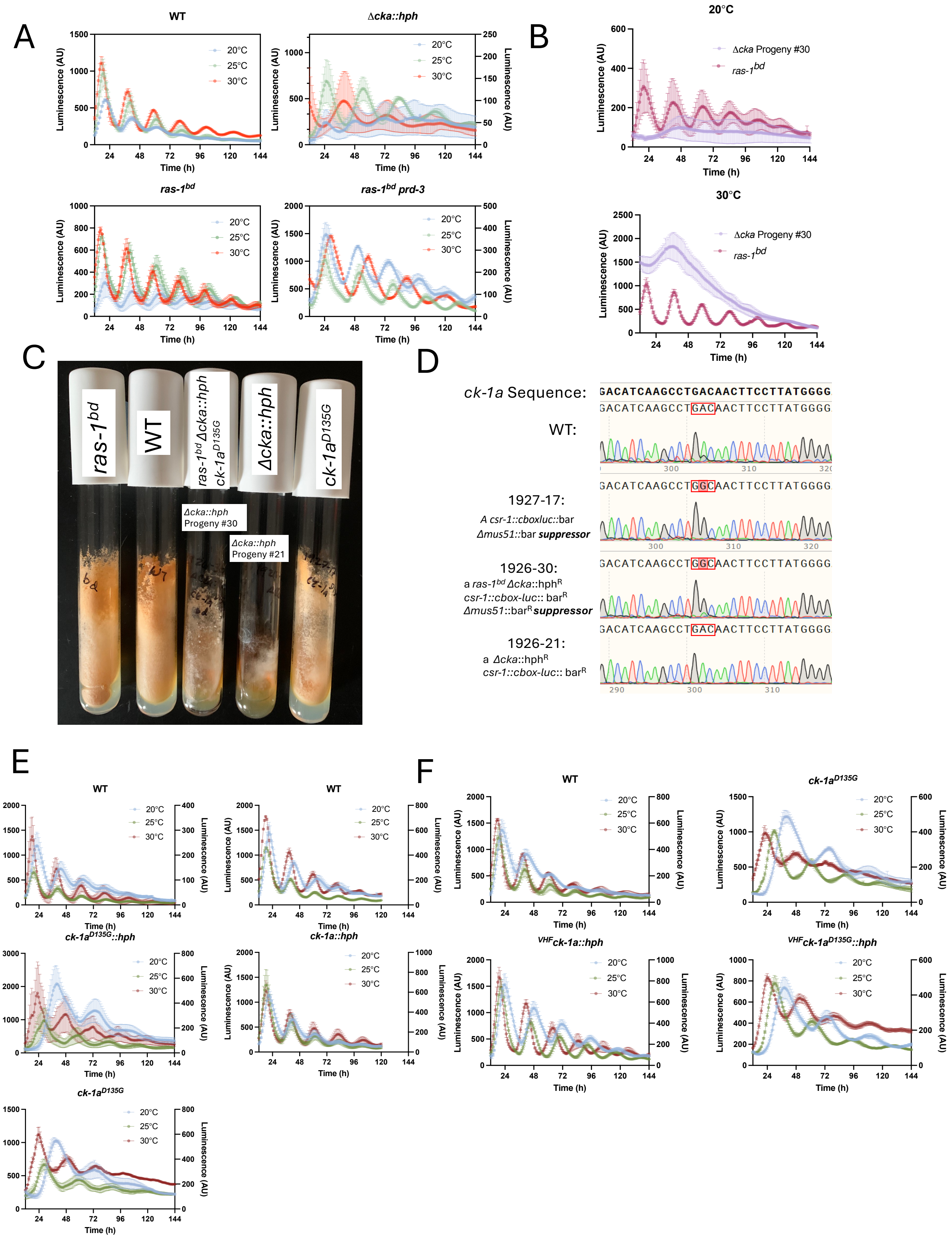
A
WT
Δcka::hph
ras-1bd
ras-1bd prd-3
Luminescence (AU)
Time (h)
20°C
25°C
30°C
B
20°C
30°C
Δcka Progeny #30
ras-1bd
C
ras-1bd
WT
ras-1bd Δcka::hph ck-1aD135G
Δcka::hph
ck-1aD135G
Δcka::hph Progeny #30
Δcka::hph Progeny #21
D
ck-1a Sequence:
GACATCAAGCCTGACAACTTCCTTATGGGG
WT:
GACATCAAGCCTGACAACTTCCTTATGGGG
1927-17:
A csr-1::cboxluc::bar
Δmus51::bar suppressor
GACATCAAGCCTGGCAACTTCCTTATGGGG
1926-30:
a ras-1bd Δcka::hphR
csr-1::cbox-luc:: barR
Δmus51::barR suppressor
GACATCAAGCCTGGCAACTTCCTTATGGGG
1926-21:
a Δcka::hphR
csr-1::cbox-luc:: barR
GACATCAAGCCTGACAACTTCCTTATGGGG
E
WT
ck-1aD135G::hph
ck-1a::hph
ck-1aD135G
F
WT
ck-1aD135G
VHFck-1a::hph
VHFck-1aD135G::hph

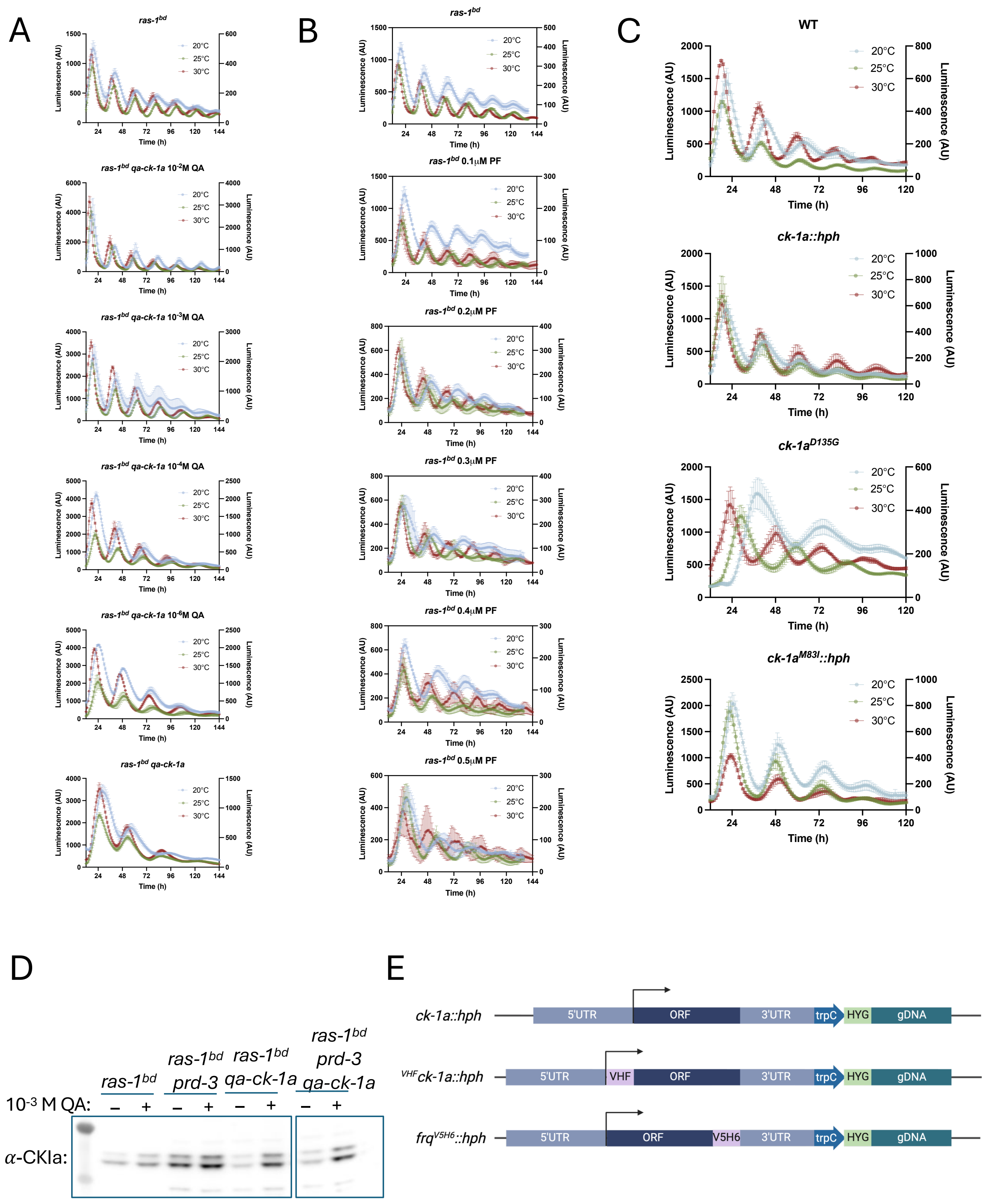
A
ras-1bd
ras-1bd qa-ck-1a 10-2M QA
ras-1bd qa-ck-1a 10-3M QA
ras-1bd qa-ck-1a 10-4M QA
ras-1bd qa-ck-1a 10-6M QA
ras-1bd qa-ck-1a
B
ras-1bd
ras-1bd 0.1μM PF
ras-1bd 0.2μM PF
ras-1bd 0.3μM PF
ras-1bd 0.4μM PF
ras-1bd 0.5μM PF
C
WT
ck-1a::hph
ck-1aD135G
ck-1aM83I::hph
20°C
25°C
30°C
Luminescence (AU)
Time (h)
D
ras-1bd
ras-1bd prd-3
ras-1bd qa-ck-1a
ras-1bd prd-3 qa-ck-1a
10-3 M QA:
α-CKIa:
E
ck-1a::hph
VHFck-1a::hph
frqV5H6::hph
5'UTR
ORF
3'UTR
trpC
HYG
gDNA
VHF
V5H6

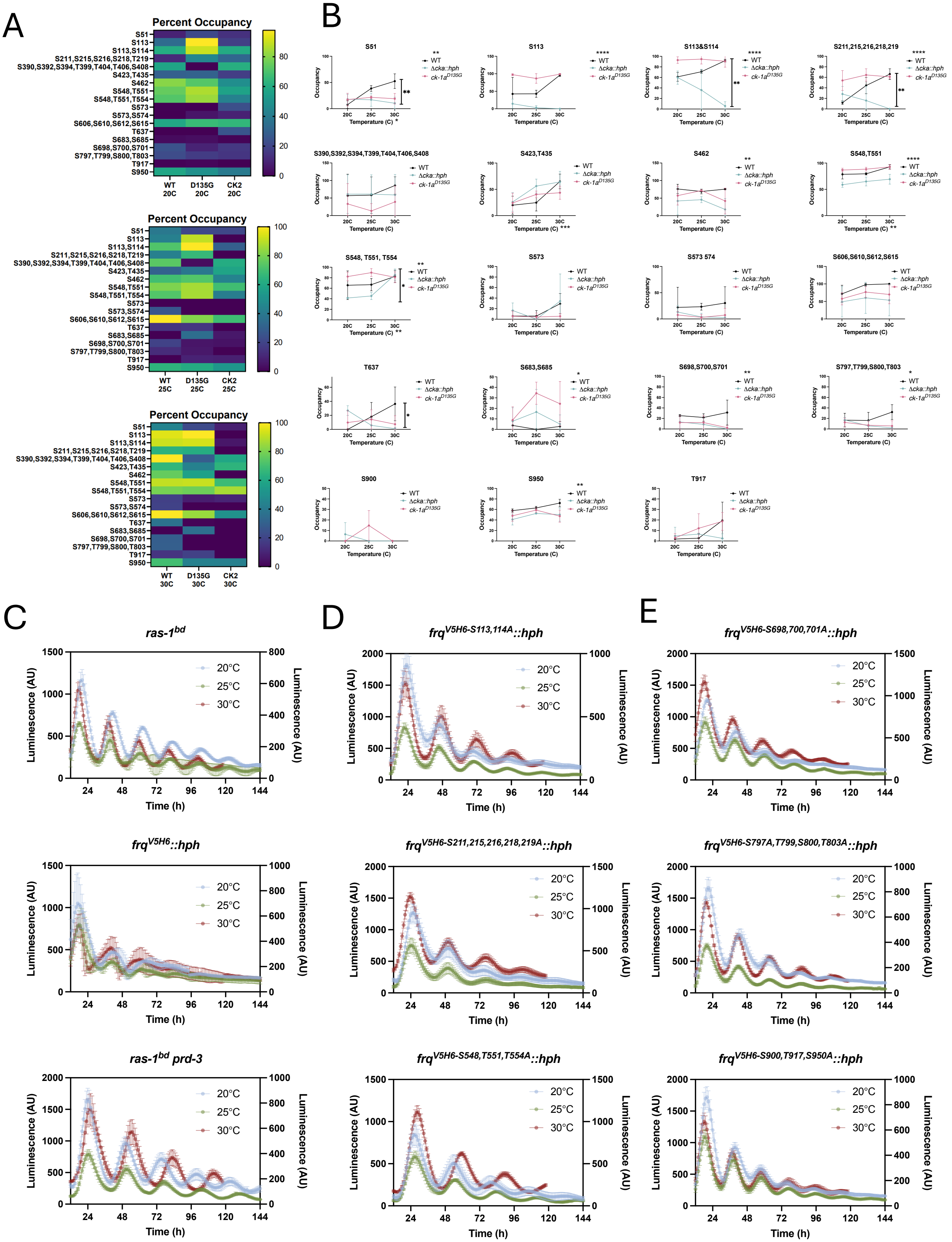
A
Percent Occupancy
S51
S113
S113,S114
S211,S215,S216,S218,T219
S390,S392,S394,T399,T404,T406,S408
S423,T435
S462
S548,T551
S548,T551,T554
S573
S573,S574
S606,S610,S612,S615
T637
S683,S685
S698,S700,S701
S797,T799,S800,T803
T917
S950
WT 20C
D135G 20C
CK2 20C
WT 25C
D135G 25C
CK2 25C
WT 30C
D135G 30C
CK2 30C
B
S51
S113
S113&S114
S211,215,216,218,219
S390,S392,S394,T399,T404,T406,S408
S423,T435
S462
S548,T551
S548, T551, T554
S573
S573 574
S606,S610,S612,S615
T637
S683,S685
S698,S700,S701
S797,T799,S800,T803
S900
S950
T917
WT
Δcka::hph
ck-1a D135G
Occupancy
Temperature (C)
20C
25C
30C
C
ras-1bd
frqV5H6::hph
ras-1bd prd-3
D
frqV5H6-S113,114A::hph
frqV5H6-S211,215,216,218,219A::hph
frqV5H6-S548,T551,T554A::hph
E
frqV5H6-S698,700,701A::hph
frqV5H6-S797A,T799,S800,T803A::hph
frqV5H6-S900,T917,S950A::hph
20°C
25°C
30°C
Luminescence (AU)
Time (h)

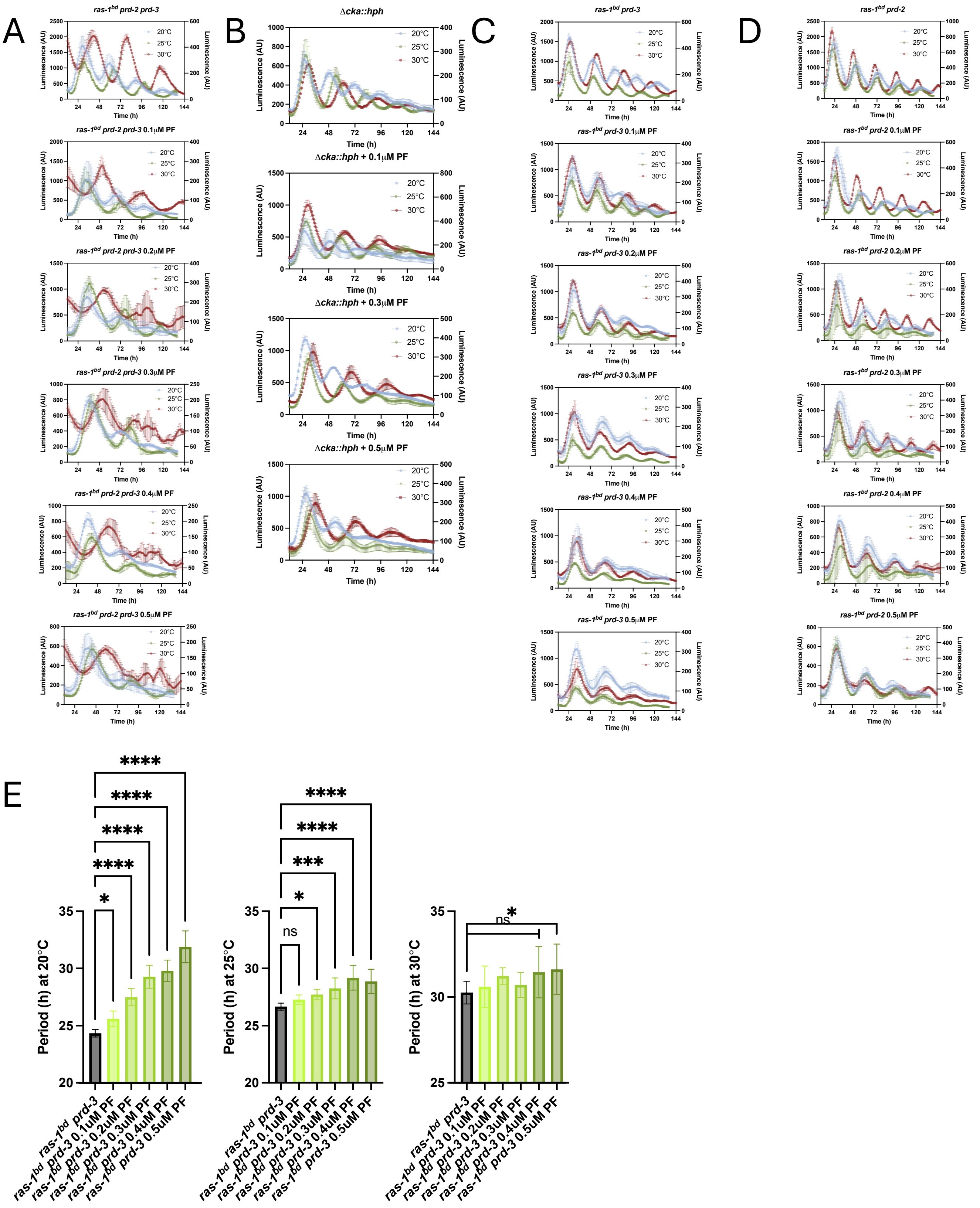

A
ras-1bd prd-2 prd-3
ras-1bd prd-2 prd-3 0.1μM PF
ras-1bd prd-2 prd-3 0.2μM PF
ras-1bd prd-2 prd-3 0.3μM PF
ras-1bd prd-2 prd-3 0.4μM PF
ras-1bd prd-2 prd-3 0.5μM PF
B
Δcka::hph
Δcka::hph + 0.1μM PF
Δcka::hph + 0.3μM PF
Δcka::hph + 0.5μM PF
C
ras-1bd prd-3
ras-1bd prd-3 0.1μM PF
ras-1bd prd-3 0.2μM PF
ras-1bd prd-3 0.3μM PF
ras-1bd prd-3 0.4μM PF
ras-1bd prd-3 0.5μM PF
D
ras-1bd prd-2
ras-1bd prd-2 0.1μM PF
ras-1bd prd-2 0.2μM PF
ras-1bd prd-2 0.3μM PF
ras-1bd prd-2 0.4μM PF
ras-1bd prd-2 0.5μM PF
20°C
25°C
30°C
Luminescence (AU)
Time (h)
E
Period (h) at 20°C
Period (h) at 25°C
Period (h) at 30°C
ras-1bd prd-3
ras-1bd prd-3 0.1uM PF
ras-1bd prd-3 0.2uM PF
ras-1bd prd-3 0.3uM PF
ras-1bd prd-3 0.4uM PF
ras-1bd prd-3 0.5uM PF
ns

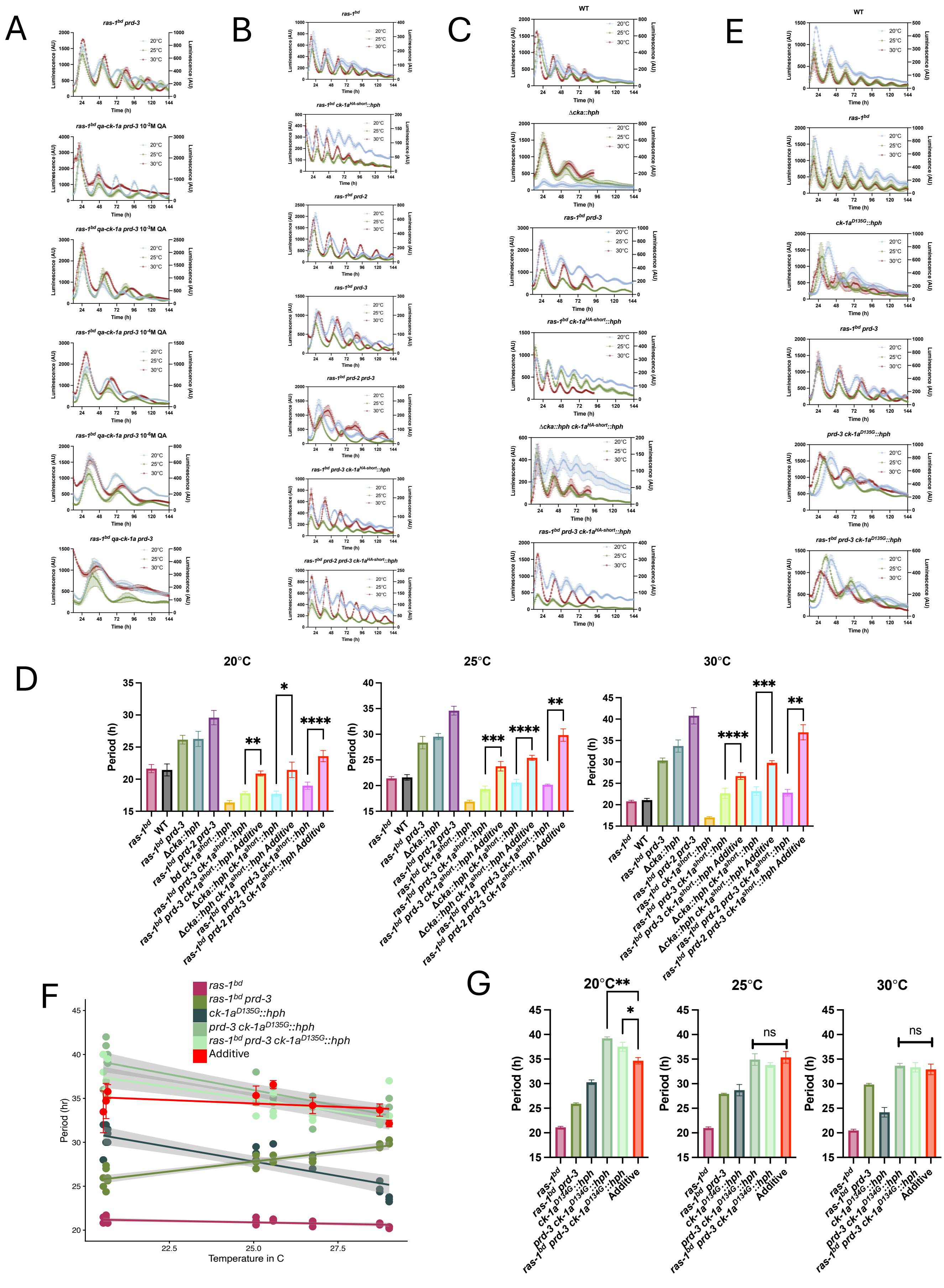
A
ras-1bd prd-3
ras-1bd qa-ck-1a prd-3 10-2M QA
ras-1bd qa-ck-1a prd-3 10-3M QA
ras-1bd qa-ck-1a prd-3 10-4M QA
ras-1bd qa-ck-1a prd-3 10-5M QA
ras-1bd qa-ck-1a prd-3
B
ras-1bd
ras-1bd ck-1aHA-short::hph
ras-1bd prd-2
ras-1bd prd-3
ras-1bd prd-2 prd-3
ras-1bd prd-3 ck-1aHA-short::hph
ras-1bd prd-2 prd-3 ck-1aHA-short::hph
C
WT
Δcka::hph
ras-1bd prd-3
ras-1bd ck-1aHA-short::hph
Δcka::hph ck-1aHA-short::hph
ras-1bd prd-3 ck-1aHA-short::hph
E
WT
ras-1bd
ck-1aD135G::hph
ras-1bd prd-3
prd-3 ck-1aD135G::hph
ras-1bd prd-3 ck-1aD135G::hph
20°C
25°C
30°C
Luminescence (AU)
Time (h)
D
20°C
25°C
30°C
Period (h)
ras-1bd
WT
ras-1bd prd-3
Δcka::hph
ras-1bd prd-2 prd-3
ras-1bd ck-1ashort::hph
ras-1bd prd-3 ck-1ashort::hph
ras-1bd prd-3 ck-1ashort::hph Additive
Δcka::hph ck-1ashort::hph
Δcka::hph ck-1ashort::hph Additive
ras-1bd prd-2 prd-3 ck-1ashort::hph
ras-1bd prd-2 prd-3 ck-1ashort::hph Additive
F
ras-1bd
ras-1bd prd-3
ck-1aD135G::hph
prd-3 ck-1aD135G::hph
ras-1bd prd-3 ck-1aD135G::hph
Additive
Period (hr)
Temperature in C
G
20°C
25°C
30°C
ns
Period (h)
ras-1bd
ras-1bd prd-3
ck-1aD134G::hph
prd-3 ck-1aD134G::hph
ras-1bd prd-3 ck-1aD134G::hph
Additive

**Supplemental Figure 1 – (A)** The average of three technical replicates luciferase traces from a 96 well plate camera run for one representative biological replicate at 20°, 25°, and 30°C for WT, *ras-1*$^{bd}$, *ras-1*$^{bd}$ *prd-3*, *Δcka::hph*. **(B)** The average of three technical replicates luciferase traces from a 96 well plate camera run for one representative biological replicate at 20°and 30°C for *bd*, and *Δcka::hph* #30, which is a sibling to *Δcka::hph* shown in A, and has the suspected suppressor mutation. **(C)** The growth of *Neurospora* strains on complete media slants in LL at 30°C for 7 days is shown for *ras-1*$^{bd}$, WT, *Δcka::hph ck-1a*$^{D135G}$ (*Δcka::hph* progeny #30, shown in B), *Δcka::hph* (*Δcka::hph* progeny #21, shown in A), and *ck-1a*$^{D135G}$ (generated from backcross of (*Δcka::hph* progeny #30 to WT). **(D)** Sanger sequencing results for the *ck-1a* locus are shown for WT, *ck-1a*$^{D135G}$, *Δcka::hph ck-1a*$^{D135G}$ (*Δcka::hph* progeny #30), *Δcka::hph* (*Δcka::hph* progeny #21). **(E)** The average of three technical replicates luciferase traces from a 96 well plate camera run for one representative biological replicate at 20°, 25°, and 30°C for WT, *ck-1a*$^{D135G}$, and *ck-1a*$^{D135G}$*::hph* run on the same plate (left), and WT and *ck-1a::hph*, run on a different plate. **(F)** The average of three technical replicates luciferase traces from a 96 well plate camera run for one representative biological replicate at 20°, 25°, and 30°C for WT, *ck-1a*$^{D135G}$, $^{VHF}$*ck-1a*$^{D135G}$*::hph,* and $^{VHF}$*ck-1a::hph*.

**Supplemental Figure 2 –** The average of three technical replicates luciferase traces from a 96 well plate camera run for one representative biological replicate at 20°, 25°, and 30°C is shown for **(A)** *ras-1*$^{bd}$, and *ras-1*$^{bd}$ *qa-ck-1a* with $10^{-2}$M, $10^{-3}$M, $10^{-4}$M, $10^{-6}$M, and 0M QA in the media; **(B)** *ras-1*$^{bd}$ with 0.1μM, 0.2μM, 0.3μM, 0.4μM, and 0.5μM PF-670462 in the media; and **(C)** WT, *ck-1a*$^{D135G}$*, ck-1a::hph, and ck-1a*$^{M83I}$*::hph*. **(D)** CKI levels were assessed with a Rabbit Anti-CKI antibody from protein extracted from *ras-1*$^{bd}$, *ras-1*$^{bd}$ *prd-3, ras-1*$^{bd}$ *qa-ck-1a,* and *ras-1*$^{bd}$ *prd-3 qa-ck-1a* grown in LCM media with and without $10^{-3}$M QA added in LL at 25°C. **(E)** Diagram of strain construction for *ck-1a::hph* strains, $^{VHF}$*ck-1a::hph* strains, and *frq*$^{V5H6}$*::hph* strains. All three constructs were targeted to the endogenous *ck-1a* or *frq* locus. A trpC promoter drives a hygromycin resistance gene. Point mutations were inserted into the constructs via Gibson Assembly from g-blocks that were synthesized with the mutation from IDT.

**Supplemental Figure 3 – (A)** A heat-map of the average phospho-occupancy on FRQ at the indicated regions across three replicates in WT, *ck-1a*$^{D135G}$, and *Δcka::hph* is shown for 20° (top), 25° (middle), and 30°C (bottom)**. (B)** A two-way ANOVA was run on all three replicates across temperature to assess statistically significant changes in phosphosite occupancy across temperature (indicated at the x-axis), between genotypes (indicated at the legend) and for the interaction between genotype and temperature (indicated within the plot) for all phosphosites that were covered in all three replicates. The average of three technical replicates luciferase traces from a 96 well plate camera run for one representative biological replicate at 20°, 25°, and 30°C is shown for **(C)** the controls – *ras-1*$^{bd}$*, ras-1*$^{bd}$ *prd-3,* and *frq*$^{V5H6}$*::hph*, **(D)** 3 phosphonull mutants that had a significant interaction effect of temperature and genotype – *frq*$^{V5H6\text{-}S113\&S114A}$*::hph***,** *frq*$^{V5H6\text{-}S211,215,216,218,219A}$*::hph*, *frq*$^{V5H6\text{-}S548,T551,T554A}$*::hph*, and **(E)** 3 phosphonull mutants that did **not** have a significant interaction effect of temperature and genotype – *frq*$^{V5H6\text{-}S698,700,701A}$*::hph, frq*$^{V5H6\text{-}S797,T799,S800,T803A}$*::hph,* and *frq*$^{V5H6\text{-}S900,T917,S950A}$*::hph*.

**Supplemental Figure 4 –** The average of three technical replicates luciferase traces from a 96 well plate camera run for one representative biological replicate at 20°, 25°, and 30°C is shown for **(A)** *ras-1*$^{bd}$ *prd-3,* **(B)** *Δcka::hph,* **(C)** *ras-1*$^{bd}$ *prd-2 prd-3*, and **(D)** *ras-1*$^{bd}$ *prd-2* with 0, 0.1μM, 0.2μM, 0.3μM, 0.4μM, and 0.5μM PF-670462 in the media**. (E)** Average period length +/- standard deviation is shown for all *ras-1*$^{bd}$ *prd-3* PF concentrations. Statistical significance was determined by a one-way ANOVA (all PF concentrations compared to *ras-1*$^{bd}$ *prd-3*).

**Supplemental Figure 5 –** The average of three technical replicates luciferase traces from a 96 well plate camera run for one representative biological replicate at 20°, 25°, and 30°C is shown for **(A)** *ras-1*$^{bd}$ *prd-3*, and *ras-1*$^{bd}$ *prd-3 qa-ck-1a* with $10^{-2}$M, $10^{-3}$M, $10^{-4}$M, $10^{-6}$M, and 0M QA in the media; **(B)** *ras-1*$^{bd}$*, ras-1*$^{bd}$ *prd-3, ras-1*$^{bd}$ *prd-2, ras-1*$^{bd}$ *prd-2 prd-3, ras-1*$^{bd}$ *ck-1a*$^{HA\text{-}short}$*::hph, ras-1*$^{bd}$ *prd-3 ck-1a*$^{HA\text{-}short}$*::hph,* and *ras-1*$^{bd}$ *prd-2 prd-3 ck-1a*$^{HA\text{-}short}$*::hph*; and **(C)** WT*, ras-1*$^{bd}$ *prd-3, Δcka::hph, ras-1*$^{bd}$ *ck-1a*$^{HA\text{-}short}$*::hph, ras-1*$^{bd}$ *prd-3 ck-1a*$^{HA\text{-}short}$*::hph,* and *Δcka::hph ck-1a*$^{HA\text{-}short}$*::hph*. **(D)** The average period length at 20°, 25°, and 30°C +/- standard deviation is shown for controls and *ras-1*$^{bd}$ *prd-3, ras-1*$^{bd}$ *prd-2 prd-3*, and *Δcka::hph* with and without *ras-1*$^{bd}$ *ck-1a*$^{HA\text{-}short}$*::hph*. Bars with bright red outline are the calculated additive period length of the sum of the period lengthening *ras-1*$^{bd}$ *prd-3, ras-1*$^{bd}$ *prd-2 prd-3,* or *Δcka::hph* contributes, plus the period shortening *bd ck-1a*$^{HA\text{-}short}$*::hph* contributes. This is the period length expected in the double mutants if these alleles act independently to regulate the clock. Statistical significance was determined by a t-test. **(E)** The average of three technical replicates luciferase traces from a 96 well plate camera run for one representative biological replicate at 20°, 25°, and 30°C is shown for WT, *ras-1*$^{bd}$*, ras-1*$^{bd}$ *prd-3, ck-1a*$^{D135G}$*::hph, prd-3 ck-1a*$^{D135G}$*::hph,* and *ras-1*$^{bd}$ *prd-3 ck-1a*$^{D135G}$*::hph*. **(F)** A linear model was fit to period lengths measured across temperature, with the grey area showing 95% confidence intervals, for *ras-1*$^{bd}$*, ras-1*$^{bd}$ *prd-3, ck-1a*$^{D135G}$*::hph*, *prd-3 ck-1a*$^{D135G}$*::hph, ras-1*$^{bd}$ *prd-3 ck-1a*$^{D135G}$*::hph,* and the calculated additive period length of the sum of period lengthening from *ras-1*$^{bd}$ *prd-3* plus period lengthening from *ck-1a*$^{D135G}$*::hph***. (G)** Average period lengths at 20°, 25°, and 30°C from (F) are shown +/- standard deviation. Statistical significance compared to Additive was determined by a one-way ANOVA (multiple comparison to additive).
